# Human ILC methylomes uncover stable lineage codes and disease-linked ILC2 regulators in allergy

**DOI:** 10.64898/2026.03.12.711290

**Authors:** Ayush Jain, Elia C. Diem, Chia-Wen Lu, Matthias Steglich, Ruth Grychtol, Maike Kosanke, Beate Pietzsch, Robert Geffers, Martin Durisin, Gesine Hansen, Anna-Maria Dittrich, Jochen Huehn, Stefan Floess, Matthias Lochner

## Abstract

The transcriptional programs of human ILCs are increasingly defined, but the DNA-methylation landscapes that stabilize their identity and function remain poorly understood. Here, we generated genome-wide DNA methylomes of human NK cells, ILC1, ILC2, and ILC3 from blood and lymphoid tissues. Subset-specific differentially methylated regions distinguished all populations and mapped to canonical regulators, including *TBX21*, *GATA3*, and *RORC*, as well as genes not previously linked to ILC biology, such as *ERN1*, *DDX47*, *JAML*, *BTLA*, and *NRROS*. Because ILC2 showed a particularly distinct methylation landscape and contribute to allergic inflammation, we tested whether selected ILC2-specific regions were functionally relevant. ILC2 marker regions were largely stable across tissues and during cytokine-driven expansion. CRISPR/Cas9-mediated deletion of the HPGDS or NRROS DMR revealed that these elements act as cis-regulatory sites controlling *HPGDS/NRROS* expression and promoting production of the type 2 cytokines IL-4, IL-5 and IL-13. Finally, methylome profiling of ILC2 from healthy, atopic, and asthmatic children identified disease-associated DMRs linked to *PTGS2*, *QKI*, and *GIMAP4*. These findings define stable epigenetic signatures of human ILC identity and uncover regulatory elements connecting ILC2 methylation to allergic disease.

## Introduction

Innate lymphoid cells (ILCs) are tissue-enriched immune cells that mirror key features of adaptive T helper cell subsets through the expression of lineage-defining transcription factors and cytokines. ILC1/NK cells, ILC2 and ILC3 are involved in barrier immunity, tissue homeostasis and inflammatory diseases, with ILC2 being particularly implicated in allergic responses and asthma.^1^ ILC subsets are commonly defined by transcription factors and effector programs: ILC1 and NK cells are linked to TBX21 and IFNγ-associated immunity,^2^ whereas ILC2 depend on GATA3 and produce type 2 cytokines such as IL-4, IL-5 and IL-13. ILC2 contribute to immune responses against parasites, but can also perform noncanonical immune cell functions, such as integrating signals from the nervous system or mediating metabolic and thermal homeostasis.^3,4^ Due to their tissue residency and capacity to express type 2 factors, ILC2 are considered key players in allergic responses.^5–7^ ILC3 are characterized by RORγt/RORC and IL-17/IL-22-associated responses, coordinating the innate immune response to both intra- and extracellular pathogens.^8^ Transcriptomic analyses of murine ILC at the single-cell level revealed considerable heterogeneity within the main ILC subpopulations found in different tissues. Combined with chromatin accessibility analyses, these results suggest the existence of tissue-specific core regulatory elements that underlie ILC identity and function.^9–11^ Similarly, scRNA-seq of ILC from various human tissues, including blood, tonsils, lungs and intestines, suggested pronounced heterogeneity and tissue-specific transcriptional imprinting.^12–16^ The combination of ATAC-seq to identify accessible chromatin and ChIP-seq to discover active promoters and enhancers led to the definition of distinct regulomes for human ILC subsets.^17,18^ These studies highlight substantial transcriptional and regulatory heterogeneity in human ILCs, but also suggest that stable epigenetic features may be particularly informative for defining lineage identity across variable tissue environments.^19^ DNA methylation is a stable epigenetic modification that contributes to immune-cell differentiation, homeostasis and effector function.^20^ In murine ILCs, genome-wide methylation profiling identified subset-specific DMRs at lineage-defining and functionally relevant loci, and methylation states were linked to transcriptional activity and regulatory element function.^21,22^ However, comparable DNA methylation maps for human ILC subsets have been lacking.

Here, we generated genome-wide DNA methylation maps of human NK cells, ILC1, ILC2 and ILC3 and identified subset-specific DMRs that define stable epigenetic marker regions. Focusing on ILC2, we show that selected marker regions are maintained during activation and expansion, nominate candidate core transcription factors associated with ILC2 DMRs, and demonstrate that *HPGDS*- and *NRROS*-linked DMRs act as functional regulatory elements controlling type 2 cytokine production. Finally, disease-resolved methylome profiling of blood-derived ILC2 from juvenile donors with atopy or asthma identified additional DMRs associated with allergic disease, including loci linked to *PTGS2*, *QKI* and *GIMAP4*.

## Results

### Human ILC subsets have distinct DNA methylation landscapes

We first asked whether human ILC subsets can be distinguished by stable DNA methylation signatures. To address this, we sorted NK cells, ILC1, ILC2 and ILC3 from human PBMCs and non-inflamed tonsils (Figures S1A-S1C). Subsequently we performed BS-seq, processed the data with the *nf-core/methylseq* analysis pipeline, and called the DMRs by pairwise comparison using the software *bsseq* (GSE313891). Principal component analysis showed tight clustering of biological triplicates and clear separation of the four populations, indicating reproducible and subset-specific methylation profiles (Figure 1A). The largest numbers of DMRs were detected between ILC2 and ILC3 (14,200) and between ILC2 and NK cells (12,345) (Figure 1B). Intermediate numbers were observed for ILC1 versus ILC2 (8,241) and ILC1 versus ILC3 (7,960), whereas ILC1 and NK cells differed least (4,912), consistent with their close relationship in hierarchical clustering (Figure 1C). Most DMRs mapped to introns (57.5-61.5%), followed by intergenic regions (22.2-24.2%). Regions spanning two annotated elements, such as promoter/intron or exon/intron combinations, were more frequent (11.1-13.9%) than DMRs confined to promoters or exons alone (< 4%), suggesting that extended regulatory segments contribute substantially to subset identity (Figure 1D). Gene Ontology analysis of DMR-associated genes linked these regions not only to general cellular programs such as transcriptional regulation, lipid metabolism and transport, but also to innate immune pathways and host-virus interactions (Figure 1E).

**Figure 1.**
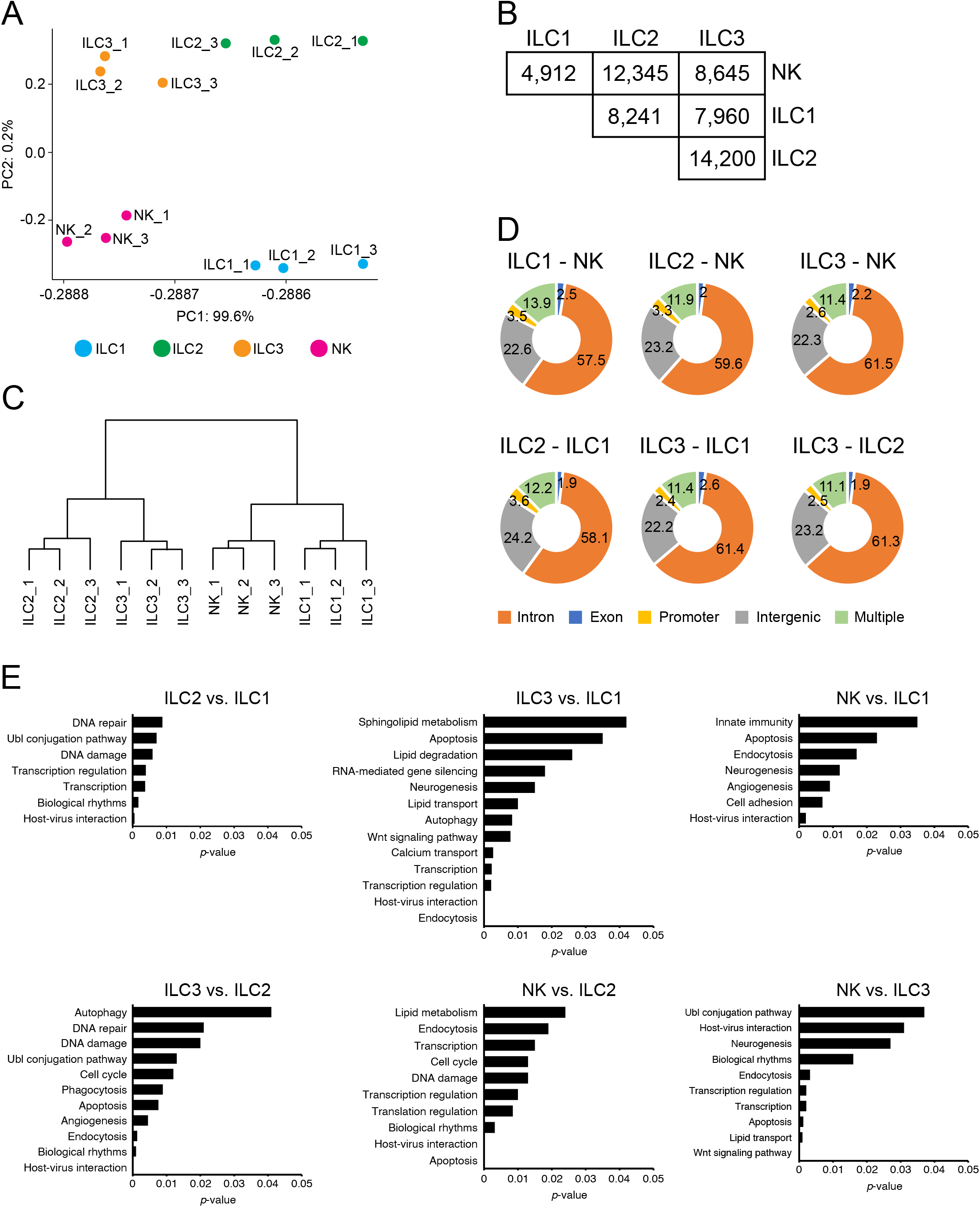
Genome-wide methylation profiling of human ILC subsets. (**A**) Principal component analysis (PCA) based on DMR mean methylation of human ILC subsets. The percentage of explained variance for each component is displayed next to the axis. (**B**) Table showing the number of DMRs identified using BSmooth from the indicated ILC methylome comparisons. (**C**) Unsupervised hierarchical clustering of the top 1,000 DMRs among ILC samples. Replicates are numbered. (**D**) Pie charts indicating the location of the DMRs identified in group-wise comparisons. Numbers indicate the frequency of DMRs in introns, exons, intergenic, promoter, or multiple regions (overlapping between 2 locations). (**E**) GO functional annotation of biological process of DMR-associated genes from pairwise methylome comparisons of ILC populations. Significance is present at *p* ≤ 0.05. Processes ranking according to *p*-value.

### Signature DMR patterns define human ILC identity across tissues

To move from global differences to lineage-defining features, we next searched for marker regions that, when combined as a signature, could uniquely identify each ILC subset with a high degree of reliability. Following criteria previously used for murine ILCs, we prioritized DMRs with > 35% methylation difference, excluded weakly differential regions and loci without clear gene annotation, and favored regions containing multiple differentially methylated CpGs that were selectively demethylated in only one subset. Using this strategy, we identified 8 to 17 of the most suitable marker regions for each ILC population, naming the regions after the nearest associated gene (Figure 2A and Table S1). As expected, many ILC1 marker regions were also demethylated in NK cells, again underscoring the close relationship between these populations, whereas most top-ranked ILC2 and ILC3 markers were subset restricted. Inspection of representative loci revealed that the marker sets recapitulated known lineage biology while also nominating unexpected candidates. In the ILC1/NK lineage, the *TBX21* locus was demethylated at the promoter in all ILCs, but demethylation extended through the first intron only in ILC1 and NK cells (Figure 2B). Similar patterns were observed at *LEF1* and *CLEC2D*, and additional ILC1-associated regions were detected at *IKZF3*, *CD6*, *CD28* and *BTLA* (Figures 2B and S2A). By contrast, a uniquely demethylated region in the *IFNG* gene was present in NK cells but not ILC1, indicating that these otherwise related populations also retain distinct methylation features (Figure S2B). In ILC2, broad demethylation was seen across the *GATA3*, *MAF* and *IL10RA* locus. Additional ILC2 marker regions were identified in genes that code for membrane-associated proteins and cytokines involved in cell migration and inflammatory responses regulation, including *KLRB1*, *KLRG1*, *ITGB1*, *HPGDS*, *NRROS*, *ERN1*, *CCR2*, *BIN3*, *IL26* and *IL4* (Figure 2C and Figure S3A). Among type 2 cytokine loci, *IL4* exhibited demethylation in ILC2, while *IL5* maintained a high degree of methylation in all ILC subpopulations and *IL13* demonstrated moderate promoter demethylation in both ILC2 and ILC3. (Figures S2B and S3A). In ILC3, demethylation centered on the *RORC* promoter and extended into intron 1 specifically in this subset, with further ILC3-specific regions identified at *KIT*, *AHR*, *IL23R* and *IL12RB1* (Figure 2D and Figure S3B). Similar to what we observed for *IL5*, the loci of the ILC3-associated cytokines *IL-22* and *IL-17A* remained largely methylated under steady-state conditions and did not contain any DMRs (Figure S2B). Together, these data show that subset-specific demethylation marks both canonical lineage-defining genes and candidate loci not previously linked to human ILC biology, including *IL26*, *ERN1*, *DDX47*, *JAML* and *BTLA*.

**Figure 2.**
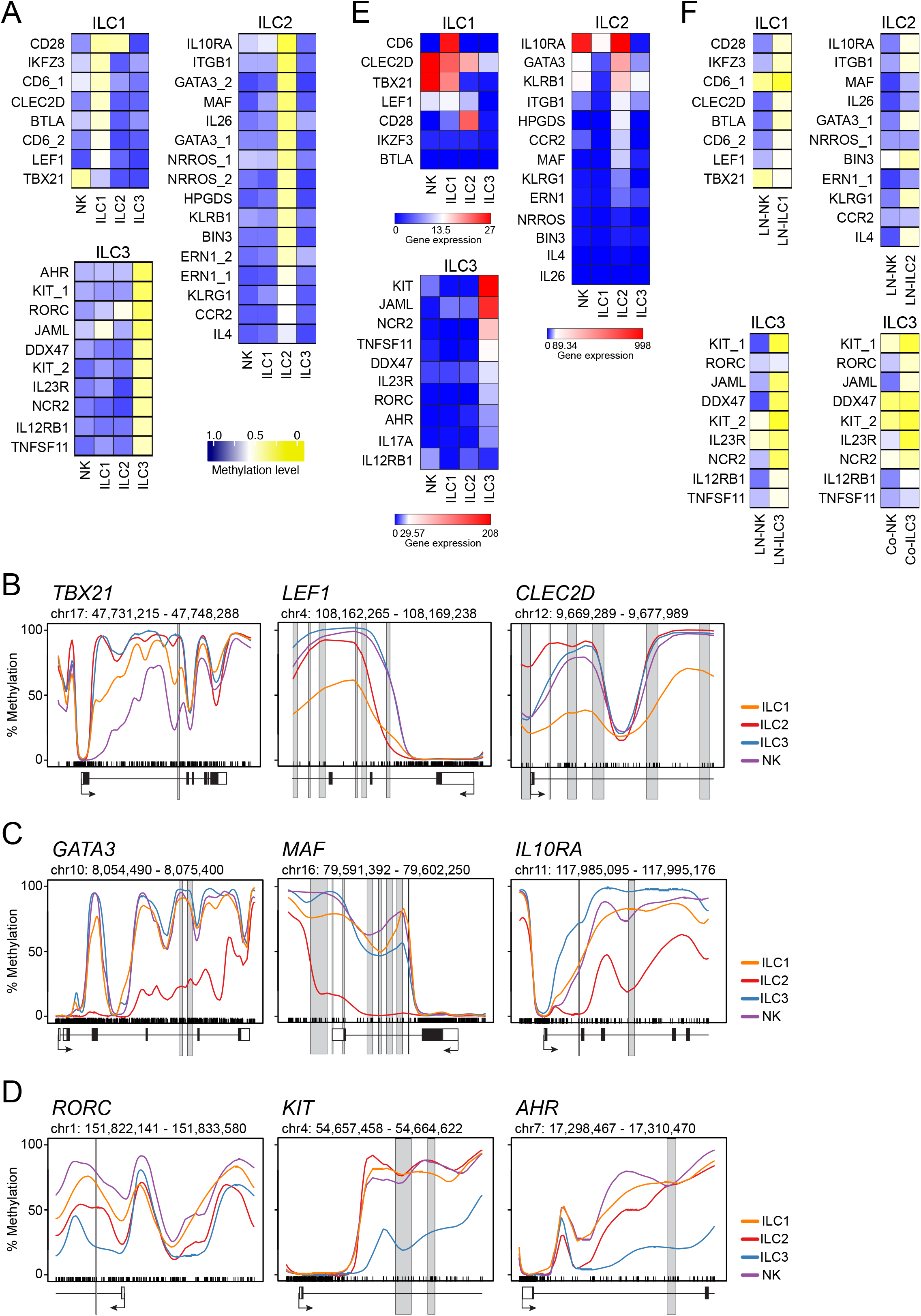
Identification of epigenetic marker regions in human ILCs. (**A**) Heat maps showing the methylation level of selected epigenetic marker regions for human ILC1, ILC2 and ILC3. The regions were named after the DMR-associated gene locus and numbered if more than one region was linked to a locus. The mean methylation values were calculated from the CpG motifs located within a marker region and translated into a color code ranging from yellow (0% methylation = 0) via white (50% methylation = 0.5) to blue (100% methylation = 1.0). (**B-D**) The methylation line plots showing genomic methylation (0-100%) as a line within the specified chromosomal location in ILC1 (orange), ILC2 (red), ILC3 (blue), and NK cells (cyan). The position of the CpG motifs is indicated by a barcode. The gene elements, transcription start site (arrow), translated (black box) and untranslated exon (white box), and the position of the DMR (grey box) are shown below or overlie the plot. Three selected gene loci for ILC1/NK (**B**), ILC2 (**C**), and ILC3 (**D**) are depicted. (**E**) Expression values (transcripts per million (TPM)) of marker-associated genes were translated into a heat map (blue-white-red) and ranked according to the respective ILC population. (**F**) Pyrosequencing of ILC subset-specific signature regions of sorted and pooled NK cells, ILC1, ILC2, and ILC3 isolated from human lymph nodes (LN, n=11) and colon (Co, n=6). The mean methylation of CpG motifs within each marker region was translated into a color code as used in (**A**).

In order to correlate the methylation status of the marker regions with the transcriptional activity of the associated genes, we analyzed previously published RNA-sequencing data from human ILCs derived from tonsil and blood.^23^ In general, expression levels of the genes correlated well with demethylation in the corresponding epigenetic region, with a few exceptions (Figure 2E). DMRs in *IL-10RA*, *KLRB1* and *ITGB1* were specifically demethylated in ILC2 but displayed broad expression in NK cells and other ILC subsets. Additionally, we observed a limited expression of *IKZF3* and *BTLA* in ILC1, as well as *IL4* and *IL26* in ILC2, despite the distinct demethylation of the epigenetic regions in these populations. (Figure 2E). In order to ascertain whether the ILC signatures were maintained across tissues, selected regions were analyzed by pyrosequencing in sorted ILC populations from human lymph nodes or NK cells and ILC3 from colonic lamina propria (Figure S4). Most ILC1 and ILC3 marker regions remained clearly demethylated in lymph-node-derived cells and retained differential methylation relative to NK cells. Several ILC2-associated regions, including *MAF*, *IL26*, *NRROS_1*, *ERN1_1* and *CCR2*, were less demethylated in lymph-node ILC2 than in blood-derived ILC2, and some regions showed demethylation in colon-derived NK cells. Overall, however, the marker panels were largely preserved across tissues, supporting the existence of robust epigenetic signatures that define human ILC identity despite tissue-dependent variation.

### ILC2 methylation signatures remain stable during activation and expansion

Because immune-cell differentiation is often accompanied by epigenetic remodeling, we next asked whether the ILC2 marker regions identified above remain stable when cells are activated and proliferate. This question is particularly relevant for ILC2, which can undergo marked expansion in response to cytokine stimulation. We therefore cultured primary human ILC2 for two weeks under type 2-promoting conditions containing IL-2, IL-7, IL-25, IL-33 and TSLP, a protocol that expanded the cells by about 100-fold while preserving expression of *CRTH2*, *CD161* and *CD127* (Figure S5). To monitor possible methylation changes during activation and proliferation, genomic DNA was collected on days 0, 7, 10 and 14 and analyzed by pyrosequencing. Most ILC2 marker DMRs remained remarkably stable throughout the culture period (Figure 3). A few regions, including *NRROS_1*, *IL10RA*, *ERN1_1* and *CCR2*, showed slightly higher methylation than in the original BS-seq dataset, whereas *NRROS_2*, *ERN1_2*, *KLRG1*, *IL4* were modestly less methylated. However, these differences were small and did not alter the overall ILC2-specific pattern. Thus, the methylation signature of human ILC2 is largely maintained during strong cytokine-driven expansion, indicating that these DMRs represent stable lineage features rather than transient activation states.

**Figure 3.**
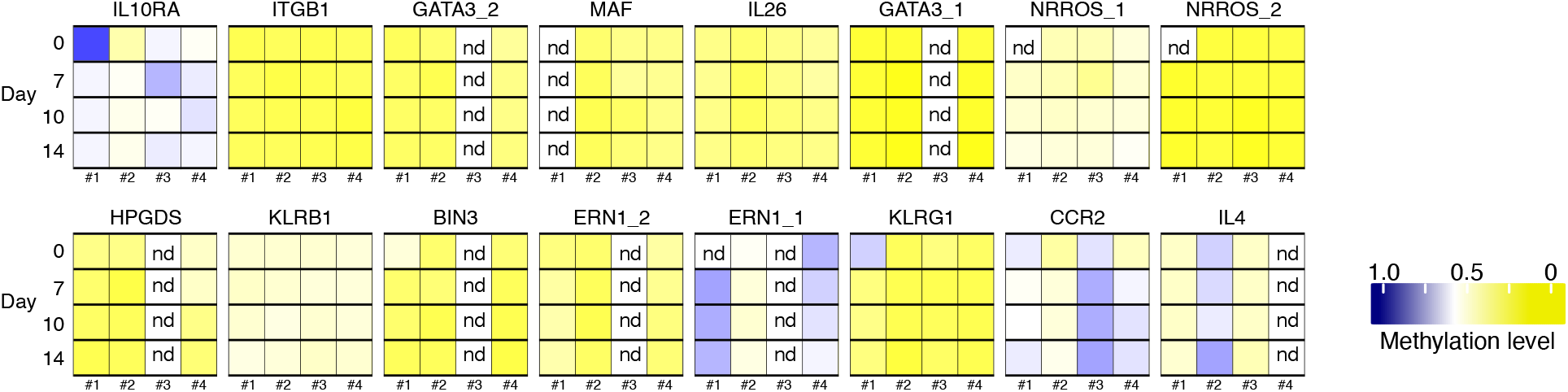
ILC2-specific marker regions show stable methylation patterns *in vitro*. Methylation levels of ILC2 marker regions in ILC2 cultured under type-II conditions *in vitro*. ILC2 from four donors (#1-#4) were harvested on days 0, 7, 10 and 14 for pyrosequencing analysis. Heat maps showing the mean methylation of the CpG motifs within each marker region, translated into a blue (100% methylation = 1), white (50% methylation = 0.5), yellow (0% methylation = 0) color code. ND, not determined.

### HPGDS and NRROS DMRs are functional regulatory elements in ILC2

To determine whether ILC2-associated DMRs merely mark lineage identity or actively regulate gene function, we used the DMRs in *HPGDS* and *NRROS* for detailed studies. In HPGDS, which encodes hematopoietic prostaglandin D2 synthase, a key enzyme required for endogenous PGD2 production in human ILC2,^24^ the DMR is located in the first intron of the gene (Figure 4A). During *in vitro* expansion, *HPGDS* expression was strongly induced under neutral (IL-2/IL-7) and especially type 2 conditions, whereas type 1 or type 3 cytokines had little effect (Figure 4B). The presence of the *HPGDS*-DMR was found to enhance the transcriptional capacity of the *HPGDS* promoter, reaching levels that surpassed those of the *CRTH2* promoter, which has been identified as active in stimulated ILC2, as evidenced by luciferase reporter assays (Figure 4C). We then disrupted the *HPGDS*-associated DMR directly in primary ILC2 using CRISPR/Cas9-mediated HDR. Deletion of this element significantly reduced *HPGDS* expression relative to control-edited cells (Figure 4D), confirming that the DMR contributes to endogenous gene regulation. Because reduced PGD2 signaling had previously been linked to impaired ILC2 activation,^24^ we also measured IL2RA and type 2 cytokines. Under standard type 2 expansion conditions, *IL2RA* expression decreased significantly after DMR targeting, whereas IL-5 and IL-13 changed only marginally (Figure 4E). We reasoned that continuous strong type 2 stimulation might mask the effect on cytokine output. Consistent with this idea, when ILC2 were first expanded under type 3 conditions, then edited, and finally restimulated for five days under type 2 conditions, DMR deletion reduced both *HPGDS* and *IL2RA* expression and significantly lowered the frequency of IL-5^+^IL-13^+^ cells (Figures 4F and 4G). These data identify the *HPGDS* DMR as a functional cis-regulatory element that supports ILC2 activation and cytokine production. We next analyzed two ILC2-associated DMRs linked to *NRROS*, one in intron 1 and one downstream of the final exon (Figure 5A). *NRROS* expression increased during ILC2 expansion, most strongly under type 2 conditions and more modestly under type 1 conditions (Figure 5B). CRISPR/Cas9-HDR-mediated deletion of either region reduced *NRROS* expression, indicating that both elements contribute to transcriptional control at this locus (Figure 5C). Because NRROS promotes degradation of NOX2,^25^ encoded by *CYBB*, we assessed *CYBB* expression after DMR targeting and found that reduced *NRROS* presence was accompanied by significantly increased *CYBB* mRNA levels (Figure 5D). Functionally, cells lacking the *NRROS*-associated regions produced significantly less IL-5 and IL-13 and showed a trend toward reduced IL-4 production compared with controls (Figures 5E and 5F). Together, these findings demonstrate that *HPGDS*- and *NRROS*-linked DMRs are not passive methylation marks but bona fide regulatory elements that shape gene expression and effector function in human ILC2.

**Figure 4.**
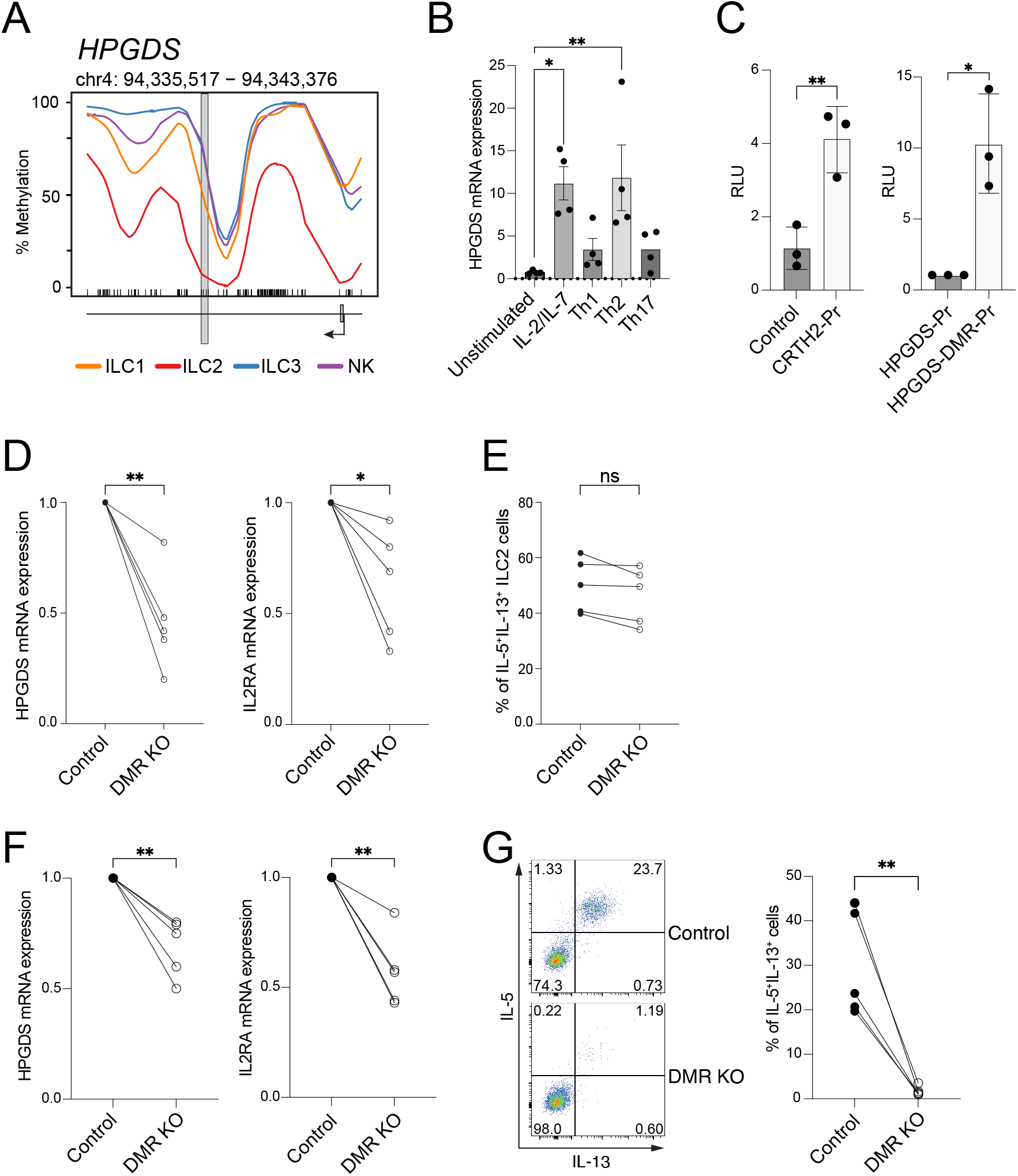
Functional analysis of *HPGDS*-associated DMR in ILC2. (**A**) Methylation line plot showing genomic methylation (0-100%) as a line within the specified chromosomal location in ILC1 (orange), ILC2 (red), ILC3 (blue), and NK cells (cyan). The position of the CpG motifs is indicated by a barcode. The gene elements transcription start site (arrow), translated (black box) and untranslated exon (white box), and the position of the DMR (grey box) are shown below or overlie the plot. (**B**) ILC2 were expanded *in vitro* in under basal (IL-2/IL-7), type I (Th1), type II (Th2) or type III (Th17) conditions. Bar graph showing expression of *HPGDS* determined by qRT-PCR before (unstimulated) and after two weeks of culture (n=4). (**C**) Bar graph showing relative light units (RLU) from luciferase reporter assays using ILC2 transfected with control plasmid (Control), *CRTH2* promoter construct (CRTH2-Pr), *HPGDS* promoter construct (HPGDS-Pr) or *HPGDS* DMR/*HPGDS* promoter constructs (HPGDS-DMR-Pr) (n=3). (**D** and **E**) *HPGDS*-associated DMR was deleted in expanded ILC2 by CRISPR/Cas9-mediated HDR (n=5). (**D**) Graph showing the expression of *HPGDS* or *IL2RA*, measured on day 6 after deletion using qRT-PCR. (**E**) In parallel, the frequency of IL-5^+^IL-13^+^ ILC2 was measured by flow cytometry. (**F**) ILC2 were expanded under type-III conditions for 14 days, followed by CRISPR/Cas9-mediated deletion of the *HPGDS*-associated DMR and 5 days of culture under type-II conditions. Expression of *HPGDS* and *IL2RA* was measured by qRT-PCR (n=5). (**G**). Representative flow cytometry plots showing IL-5 and IL-13 positive cells in expanded ILC2 following the CRISPR/Cas9-mediated deletion of the *HPGDS*-associated DMR (bottom) or the HsAAV1 control (top). Graph depicting the frequency of IL-5^+^IL-13^+^ cells obtained from all experiments (right, n=5). One-way ANOVA with Dunnett’s multiple comparison (**B**) or Paired t-test (**C**-**G**). p > 0.05: not significant (ns), p ≤ 0.05: *, p ≤ 0.01: **, p ≤ 0.001: ***.

**Figure 5.**
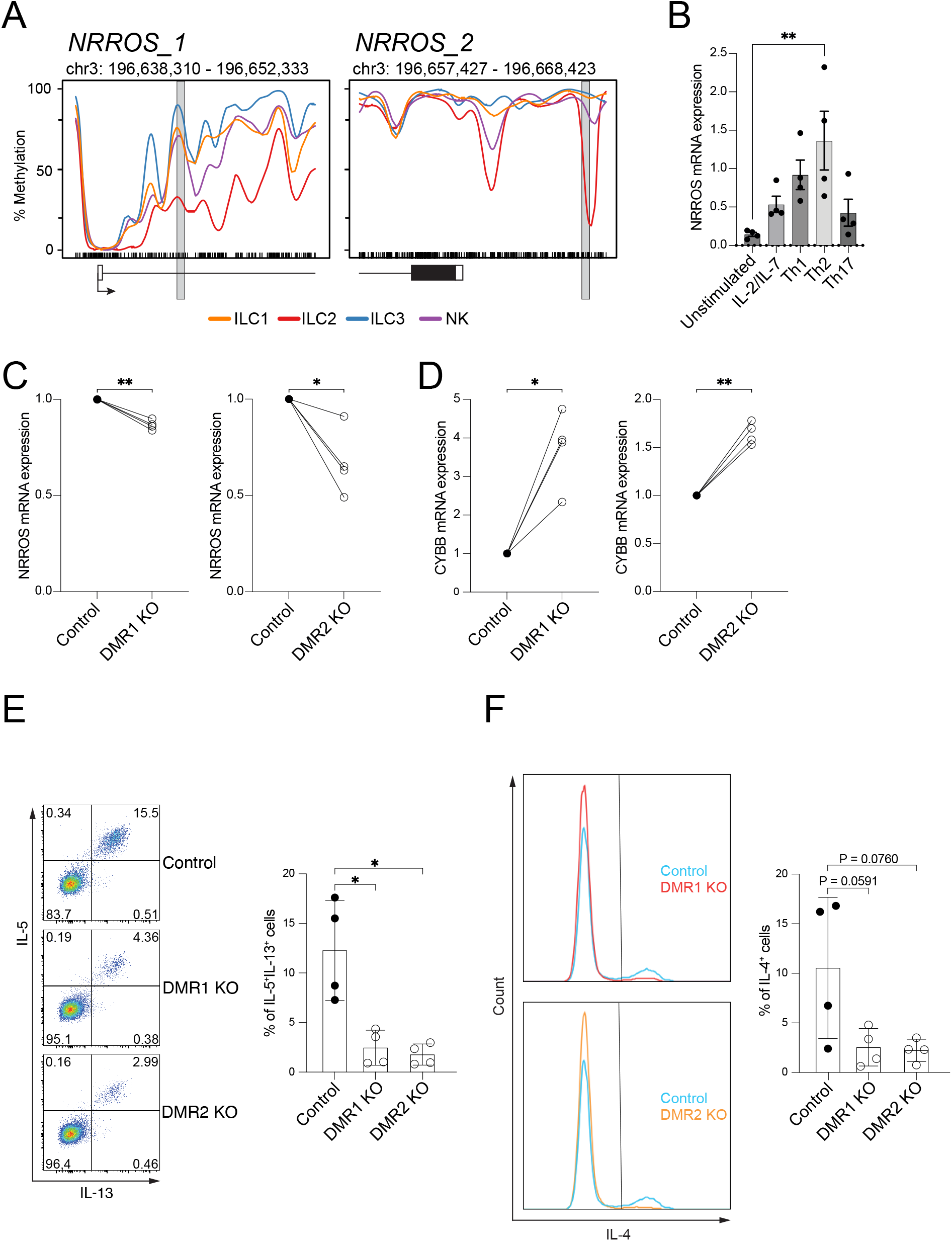
Functional analysis of NRROS-associated DMR in ILC2. (**A**) Methylation line plots showing genomic methylation (0-100%) as a line within two locations in *NRROS* in ILC1 (orange), ILC2 (red), ILC3 (blue), and NK cells (cyan). The position of the CpG motifs is indicated by a barcode. The gene elements transcription start site (arrow), translated (black box) and untranslated exon (white box), and the position of the DMR (grey box) are shown below or overlie the plot. (**B**) ILC2 were expanded under basal (IL-2/IL-7), type-I (Th1), type-II (Th2) or type-III (Th17) conditions. Bar graph showing expression of *NRROS* determined by qRT-PCR before (unstimulated) and after two weeks of culture (n=4). (**C** and **D**) NRROS-associated DMRs (DMR1 KO, DMR2 KO) were deleted in expanded ILC2 by CRISPR/Cas9-mediated HDR. Graph showing the expression of *NRROS* (**C**) and *CYBB* (**D**) on day 5 after deletion using qRT-PCR (n=4). (**E** and **F**) Representative flow cytometry analysis showing the expression of IL-5 and IL-13 (**E**) or IL-4 (**F**) in expanded ILC2 with CRISPR/Cas9-mediated deletion of the NRROS-associated DMRs (DMR1 KO, DMR2 KO) or HsAAV1 (Control) (n=4). One-way ANOVA (repeated measures) with Dunnett’s multiple comparison (**B, E** and **F**) or Paired t-test (**C** and **D**). p > 0.05: not significant (ns), p ≤ 0.05: *, p ≤ 0.01: **, p ≤ 0.001: ***.

### ILC2 DMRs share a candidate core transcription-factor network

The functional activity of *HPGDS-* and *NRROS*-linked DMRs suggested that ILC2-associated DMRs may be coordinated by a broader regulatory network. We therefore asked whether these regions share enriched transcription-factor motifs that could help explain their common activity. To address this, we performed motif enrichment analysis across all ILC2-associated DMRs identified in pairwise comparisons with ILC1, ILC3 and NK cells (Figure 6A). Binding motifs for 51 transcription factors were enriched in the ILC2-versus-ILC1 comparison, 35 in ILC2-versus-ILC3 and 46 in ILC2-versus-NK, after restricting the analysis to factors expressed in ILC2 according to published RNA-seq data^23^ (Figures 6A and 6B). The enriched factors included KLF family members associated with DNA demethylation, immune-related regulators such as STAT2 and IRF family proteins, and several zinc-finger proteins including ZNF770, ZNF263 and ZNF121. Additional candidates included MAZ, PATZ1, FOXJ2 and LEF1. Notably, 25 transcription factors were shared across all three comparisons, suggesting the existence of a core ILC2-associated regulatory module (Figure 6B). Within the *NRROS_1* DMR we identified predicted binding sites for KLF4, IRF1/3, STAT2 and ZNF121; *NRROS_2* contained motifs for ZNF770, ZNF394 and ZNF121; and the *HPGDS* DMR also harbored predicted ZNF770 and ZNF121 sites (Figures S6A and S6B). These analyses define a common set of candidate transcription factors that may connect DNA methylation state to transcriptional activation and effector programming in ILC2.

**Figure 6.**
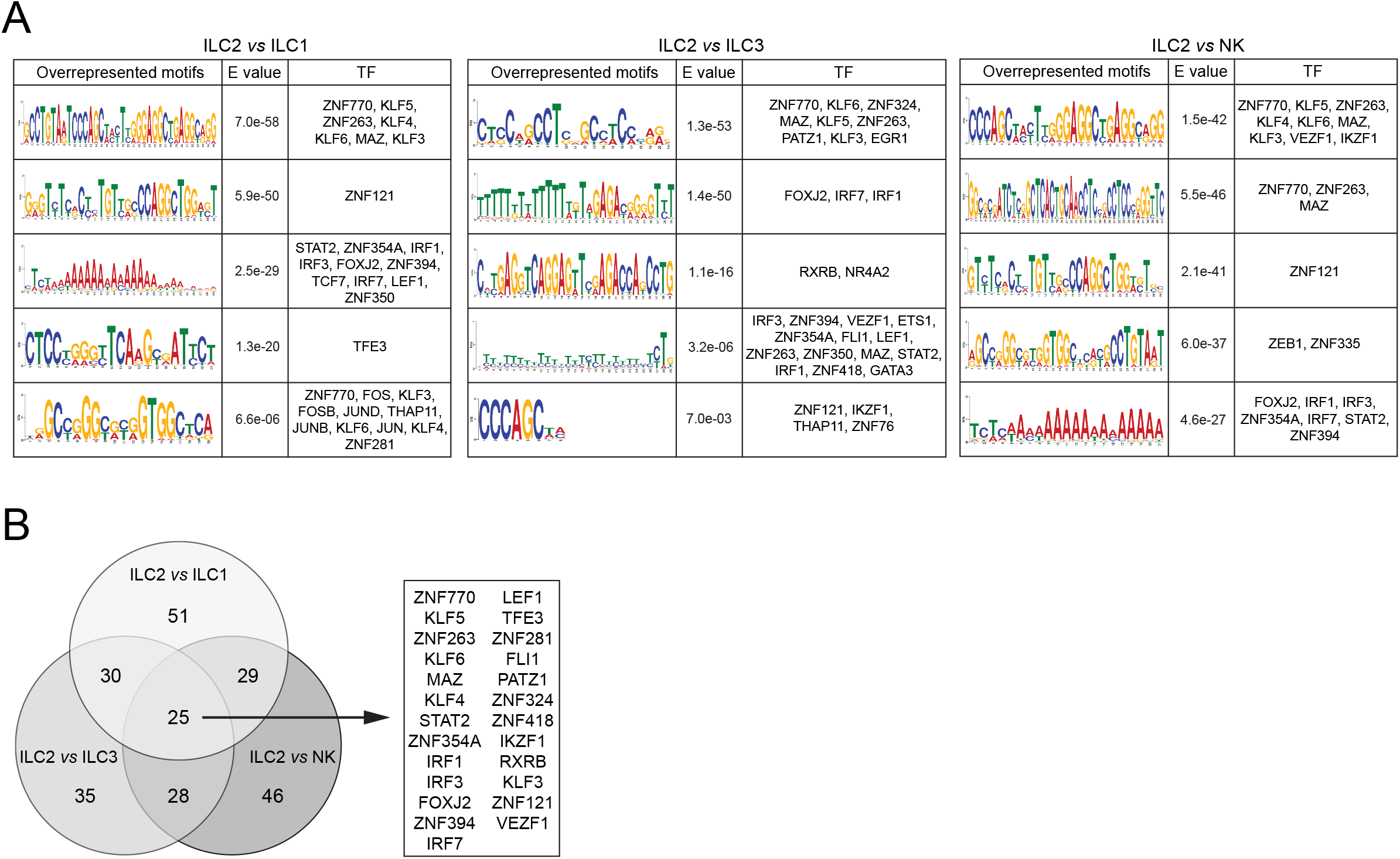
Overrepresented motifs in ILC2 DMRs define epigenetically linked core TFs for ILC2. (**A**) Table depicting the top 5 overrepresented sequence motifs and corresponding E values determined by MEME analysis, including all identified ILC2 DMRs. A search for transcription factor binding sites identified putatively associated TFs. The right columns show TFs expressed in ILC2. (**B**). Venn diagram showing the number of common transcription factors associated with DMRs identified in different pairwise comparisons. ILC2 core transcription factors within the overlap of all three comparisons are listed on the right.

### Pediatric atopy and asthma are associated with additional methylation changes in ILC2

Having defined stable ILC2 marker regions in healthy adults, we finally asked whether these signatures are preserved in pediatric disease and whether atopy or asthma introduce additional methylation changes. We therefore analyzed blood-derived ILC2 from juvenile healthy donors and juvenile patients with atopy or asthma. Because ethically permissible blood volumes from pediatric donors limited the number of directly obtainable ILC2, cells were expanded prior to genome-wide methylation analysis by EM-seq (Figure S7 and Table S2). We first examined the ILC2-specific marker regions identified in the adult discovery cohort. With the exception of *CCR2*, these regions showed comparable demethylation in healthy, atopic and asthmatic donors, indicating that the core ILC2 methylation signature is largely stable across age and clinical status (Figure 7A; GSE314735). By contrast, *IL13* and *IL5* were more strongly demethylated in all juvenile samples than in freshly isolated adult ILC2, most likely reflecting the *ex vivo* type 2 expansion required before sequencing, whereas non-type 2 loci such as *TBX21* and *IL23R* remained unchanged (Figures S2B, 3B and S8A).

**Figure 7.**
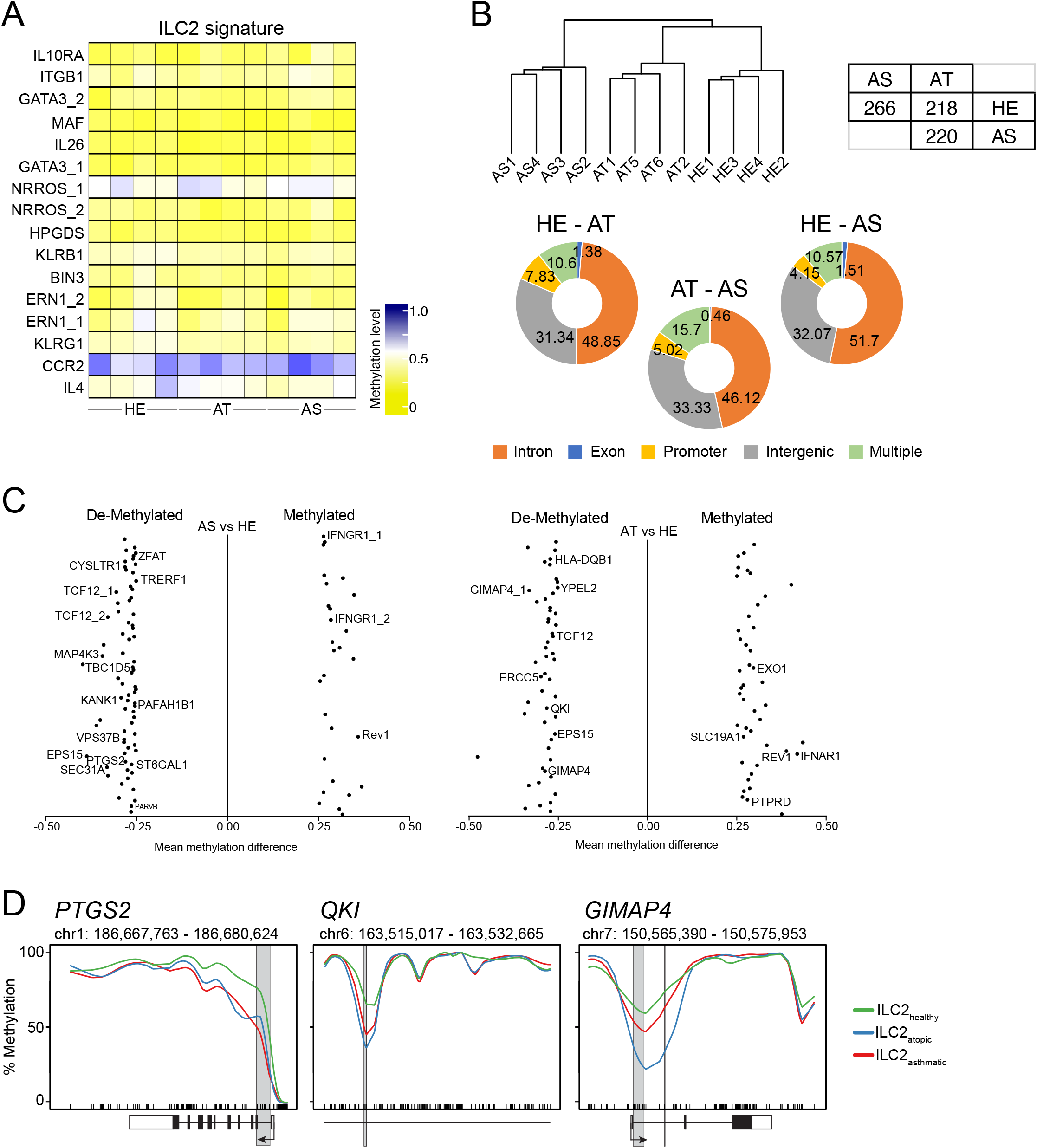
Genome-wide methylation profiling of human ILC2 from atopic and asthmatic patients. (**A**) Heat map showing the methylation level of ILC2 marker regions in expanded ILC2 derived from peripheral blood of asthmatic, atopic, and healthy donors. The mean methylation values were calculated from the CpG motifs located within a marker region and translated into a color code ranging from yellow (0% methylation = 0) via white (50% methylation = 0.5) to blue (100% methylation = 1.0). (**B**) Unsupervised hierarchical clustering of the top 1,000 ILC2 DMRs among patient samples; healthy controls (HE; n=4), atopic (AT; n=4) and asthma (AS; n=4). Replicates are numbered. Table showing the number of DMRs among ILC2 populations in pairwise comparisons. Pie charts depicting the location of the DMRs identified in group-wise comparisons. Numbers indicate the frequency of DMRs in introns, exons, intergenic, promoter or multiple regions (overlapping between 2 locations). (**C**) Volcano plot of identified DMRs in AS vs HE and AT vs HE comparisons. DMRs with > 25% methylation difference that could be allocated to annotated gene loci were included. (**D**) Methylation line plots showing genomic methylation (0-100%) as a line within the specified chromosomal location in ILC2 from healthy (green), atopic (blue), and asthmatic (red) donors. The position of the CpG motifs is indicated by a barcode. The gene elements transcription start site (arrow), translated (black box) and untranslated exon (white box), and the position of the DMR (grey box) are shown below or overlie the plot.

Despite the preservation of baseline ILC2 markers, unsupervised analysis revealed disease-associated methylation differences beyond the core signature. Hierarchical clustering based on ILC2-associated DMRs grouped samples according to donor category and clearly separated healthy, atopic and asthmatic replicates (Figure 7B). Pairwise comparisons identified 220 DMRs between asthmatic and atopic donors, 266 between asthmatic and healthy donors, and 218 between atopic and healthy donors, substantially fewer than observed between major ILC lineages but sufficient to distinguish disease states (Figure 7B). As in the adult dataset, most DMRs mapped to introns (48.85-51.7%) and intergenic regions (31.34-33.33%), and overlaps involving promoters together with introns or exons (10.5-15.7%) were more common than promoter-only or exon-only regions (<= 5%).

To gain insight into disease-driven mechanisms, we performed Gene Ontology (GO) analyses using DMR-associated genes from the asthmatic versus healthy and atopic versus healthy comparisons (Figure S8B). In the asthmatic versus healthy comparison, we identified genes associated with asthma and inflammation, including *IFNGR1*, *PTGS2*, *KANK1*, *CYSLTR1*, *PARVB*, and *PAFAH1B1* (Figure 7C). Genes annotated to tobacco use disorder (*MAP4K3*, *TCF12*, *TRERF1*, *ZFAT*, *TBC1D5*, *ST6GAL1*, *VPS37B*) showed a trend toward enrichment, consistent with the known exacerbating role of smoking in asthma. Similarly, GO analysis of DMR-associated genes from the atopic versus healthy comparison revealed genes associated with lung dysfunction and allergy, including *EPS15*, *HLA-DQB1*, *QKI*, *PTPRD*, *YPEL2*, *REV1*, *IFNAR1*, *SLC19A1*, *ABCC1*, *EXO1*, and *ERCC5* (Figure 7C). Methylation profiles of selected loci revealed pronounced demethylation around DMRs linked to *PTGS2*, *QKI* and *GIMAP4* in ILC2 from atopic and asthmatic donors, nominating these loci as candidate disease-associated regulatory regions (Figure 7D). Together, these data show that the core ILC2 methylation signature is preserved in juvenile donors and allergic disease, while disease-resolved methylome profiling reveals additional DMRs that distinguish atopic and asthmatic ILC2 from healthy controls. This expanded marker set links human ILC2 methylation programs to allergic disease and nominates candidate regulatory loci for future functional studies.

## Discussion

Epigenetic mechanisms play an important role in immunology, as they affect key processes such as the development, differentiation, and function of various immune cell types.^20^ While epigenetic regulation in adaptive immunity has been extensively studied, research on innate immune cell types like innate lymphoid cells (ILCs) is limited, particularly in humans. Previous studies in this area have primarily focused on histone modifications, chromatin accessibility, and long non-coding RNAs due to their technical feasibility,^26^ and a global analysis of DNA methylation in ILCs has only been performed in mice so far.^21,22,27,28^ To address this knowledge gap, this study performed genome-wide methylation sequencing of human ILC subsets, providing CpG methylation datasets at single CpG resolution.

The genome-wide methylation analysis of ILC1, ILC2, ILC3 and NK cells enabled us to identify a set of DMRs that are specific to the various ILC subsets in humans. In line with studies in mice, we found substantial demethylation within the first intron of the *TBX21* locus in both ILC1 and NK cells, highlighting the role of this transcription factor in the function and stability of these populations. Despite the well-described molecular and functional similarities between ILC1 and NK cells, which were also reflected by the relatively small number of DMRs identified in our analysis, we were able to pinpoint regions that were preferentially demethylated in ILC1. These included regions in the *CD6* locus, consistent with reported *CD6* expression in human ILC1,^12,23^ as well as a marker region in *LEF1*, a Tcf/Lef family transcription factor involved in lymphoid development and Wnt signaling.^29–31^ While the precise functions of CD6 and LEF1 in human ILC1 remain to be defined, their gene loci illustrate how methylation profiling can nominate candidate regulators beyond canonical lineage factors. Unlike murine ILCs, where we did not identify marker regions in the locus of the ILC3- and LTi-associated transcription factor RORγt (encoded by *RORC*), we found a demethylated region in the *RORC* gene locus in human ILC3. This is consistent with previous observations in human Th17 cells and RORγt-expressing regulatory T cells,^32^ suggesting that epigenetic mechanisms contribute to the regulation of RORγt expression in humans. In addition to regions linked to established ILC3-associated genes such as *AHR*, *KIT*, *IL23R*, *TNFSF11* and *NCR2*, we identified ILC3-specific regions in *JAM*L and *DDX47*. JAML has been associated with leukocyte transmigration and tissue immunity,^33^ whereas DDX47 has mainly been studied in the context of genome stability and cancer.^34,35^ Their identification as ILC3-associated marker loci therefore points to candidate regulatory programs that remain to be explored in human ILC3. Regarding ILC2, the identification of DMRs in *GATA3* was expected, given its central role in ILC2 development and function. Notably, the two DMRs identified in our analysis are located in functional enhancer elements previously implicated in ILC2 biology.^36^ In addition, we observed extensive demethylation of the *MAF* locus, consistent with the role of MAF in regulating pro- and anti-inflammatory cytokine programs in ILC2.^37,38^ We also identified DMRs in loci encoding prototypic human ILC2-associated molecules such as *KLRB1* and *KLRG*1, as well as in *IL4* and *IL26*. Although *IL4* and *IL26* are not strongly expressed under steady-state conditions, their demethylation may indicate regulatory potential that becomes relevant upon activation. Together, these findings indicate that the human ILC marker regions identified here capture established lineage-defining programs while also highlighting less explored candidate loci for future functional studies.

HPGDS is a key enzyme responsible for intrinsic PGD2 production, which is essential for human ILC2 activation.^24^ Our functional analysis of the DMR located within the *HPGDS* locus revealed a regulatory role for this epigenetic region, as CRISPR/Cas9-mediated deletion reduced *HPGDS* expression, impaired *IL2RA* induction and decreased IL-5/IL-13 production. Thus, the HPGDS-associated marker region is not only indicative of ILC2 identity, but contributes to the regulation of an activation pathway relevant for ILC2 effector function. We further identified two DMRs in the *NRROS* locus that mediated transcriptional activity. Deletion of these regions reduced *NRROS* expression and resulted in decreased IL-4, IL-5 and IL-13 production, suggesting that NRROS positively contributes to type 2 cytokine production in human ILC2. Since NRROS inhibits reactive oxygen species production through proteasomal degradation of NOX2,^25^ we also examined *NOX2*/*CYBB* expression and found it to be increased in *NRROS* DMR-targeted cells. Although further experiments are required to define the underlying mechanism, these findings nominate NRROS as a previously unrecognized regulator of human ILC2 function. Collectively, the DMR deletion approach in *HPGDS* and *NRROS* demonstrates that selected ILC2-associated DMRs are not merely epigenetic markers, but functional cis-regulatory elements that modulate gene expression and effector cytokine production.

The methylome datasets obtained from healthy adult donors define core epigenetic signatures of human ILC subsets, but may not capture age- or disease-associated regulatory changes. Therefore, we extended our analysis to ILC2 isolated from juvenile healthy, atopic and asthmatic donors and identified over 200 DMRs in each pairwise comparison between donor groups. Among regions demethylated in both asthmatic and atopic ILC2, we identified a DMR within the first intron of *PTGS2*, encoding cyclooxygenase-2 (COX-2), a key enzyme in prostaglandin synthesis. This finding is notable because COX-2 inhibition has been shown to impair IL-33-mediated proliferation and IL-5/IL-13 production in ILC2, and *PTGS2* polymorphisms have been associated with pediatric asthma, atopy and bronchitis.^39–41^ Another region demethylated in both asthmatic and atopic ILC2 was located in the *QKI* locus. QKI proteins regulate several aspects of RNA metabolism, including splicing, mRNA stability and transport, and have been linked to regulation of Notch signaling through alternative splicing of *NUMB*.^42^ Since Notch signaling modulates allergic inflammation,^43^ epigenetic regulation of *QKI* may contribute to altered ILC2 function in allergic disease. Among regions specifically demethylated in atopic patients, we identified two DMRs in the first intron of *GIMAP4*, a GTPase-family gene implicated in lymphocyte function and previously associated with allergic sensitization and asthma in children.^44–46^ Although the role of GIMAP4 in ILCs remains unknown, these findings nominate *GIMAP4* as a candidate regulatory locus in allergy-associated ILC2 responses.

Together, the disease-resolved methylome analysis shows that the core ILC2 methylation signature is preserved across donor groups, while additional DMRs distinguish ILC2 from atopic and asthmatic donors. These disease-associated marker regions provide a first methylation-based framework for studying human ILC2 in allergic disease and nominate *PTGS2*, *QKI* and *GIMAP4* as candidate loci for future functional validation.

## Limitations of the study

Our primary methylome maps were generated from bulk-sorted ILC subsets isolated from different anatomical sources, including circulating ILC2 from blood and ILC1/ILC3 from tonsils. Although analyses of lymph-node and colonic ILCs support the broad conservation of selected marker regions, tissue-resident populations may exhibit additional niche- or state-dependent methylation patterns. Future studies using single-cell methylome profiling combined with matched transcriptomic and spatial analyses will be required to resolve intra-subset heterogeneity and tissue specificity more precisely. The disease-resolved analyses in juvenile donors relied on peripheral-blood ILC2 and required an *ex vivo* expansion step to obtain sufficient cell numbers for EM-seq, as the ethically permissible blood volumes from pediatric donors were limited and precluded direct methylome profiling without culture. Patient cohort sizes were modest, limiting statistical power and potentially introducing residual confounding despite standardized workflows. Finally, functional validation focused on two ILC2 loci (*HPGDS* and *NRROS*) and was performed *in vitro*. Whether analogous regulatory mechanisms operate in tissue ILCs *in vivo* remains to be tested.

## Data availability

- Sequencing data and the DMR result tables from the pairwise comparisons is available via GEO: GSE313891 for the healthy, adult donors and GSE314735 for the healthy, atopic and asthmatic young donors.

## Acknowledgments

We thank Lothar Gröbe for performing flow cytometry-based cell sorting. We acknowledge the assistance of the Genome Analytics Facility of the Helmholtz Centre for Infection Research and the Cell Sorting Core Facility of the Hannover Medical School, supported in part by Braukmann-Wittenberg-Herz-Stiftung and Deutsche Forschungsgemeinschaft. This work was supported by grants from the German Research Foundation within the priority program 1937 on innate lymphoid cells (DFG LO 1415/8-2 to M. Lochner, DFG FL 965/1-2 to S. Floess, and DFG DI 1224/6-1 to AM. Dittrich) and the German Federal Ministry for Research, Technology and Space (BMFTR) via a grant for the German Center for Lung Research (DZL; FKZ 82DZL002C1) to AM. Dittrich.

## Author contributions

Author contributions: AJ, ECD, CL and BP performed the experiments. MS and MK established and performed all methylome analyses. RoG reanalyzed published RNA-seq data. RuG, MD, GH, AMD and JH provided essential materials and helped with interpretation of the data. SF and ML designed the study, interpreted the data, and together with AJ wrote the manuscript.

## Material and methods

### Human samples and ethics statement

Non-inflamed human tonsils and lymph nodes were provided by the Department of Otorhinolaryngology at Hannover Medical School (MHH). Peripheral human blood from healthy donors was delivered by the Institute of Transfusion Medicine and Transplant Engineering, MHH. Non-inflamed human colon tissue was provided by the Department of Gastroenterology, Hepatology, Infectious Diseases and Endocrinology, MHH. Pediatric blood samples were collected in the Department for Pediatric Pneumology, Allergology and Neonatology of the MHH from male patients with asthma and healthy controls (Table S2). Asthma was diagnosed in accordance with the Global Initiative for Asthma (GINA) and national guidelines.^47^ Atopy was defined by the presence of allergen-specific IgE ≥ 0.7 kU/L to at least one allergen from a panel comprising 36 food and inhalant allergens (Euroline™, Euroimmun). Donors without allergen-specific IgE levels below 0.35 kU/L were considered as healthy. Informed consent from all donors was obtained. All works with human cells were approved by the ethical committee of the MHH under permit numbers: 3082-2016, 8676_BO-K_2019, 3639–2017, 10292_BO_K_2023, and 1672-2013.

### Antibodies

The following antibodies were purchased from BioLegend: CD1a-APC (HI149), CD3-APC (OKT3), CD3-Pacific Blue (OKT3), CD19-APC (SJ25C1), TCRαβ-APC (IP26), TCRγδ-APC (B1), CD14-APC (63D3), CD11c-APC (Bu15), FcεRIα-APC (AER-37), CD45-AF488 (HI30), CD127-BV421 (A019D5), CD56-APC-Fire750 (5.1H11), CD159a (NKG2A)-PE (S19004C), CD117-PE-Cy7 (104D2), CD294 (CRTH2)-BV605 (BM16), CD336 (NKp44)-PerCP-Cy5.5 (P44-8), CD4-BV711 (RPA-T4), CD25-PE (BC96), CD161-PE/Dazzle594 (HP-3G10), CD45RA-BUV805 (HI100), CCR6-BV421 (G034E3), CCR7-PE/Cy7 (G043H7), CXCR3-AF488 (G025H7), IL-5-PE (JES1-39D10), IL-4-PerCP/Cy5.5 or IL-4-PE/Dazzle594 (MP4-25D2) and IL-13-PE-Cy7 (JES10-5A2. NKG2A-PE (Z199) was purchased from Beckman Coulter and CD127-BV421 (HIL-7R-M21) from BD Biosciences.

### Cell isolation and flow cytometry

Human peripheral blood mononuclear cells (PBMCs) from either whole blood samples or leukocyte reduction systems were first enriched by using a Lymphoprep gradient in combination with a SepMate-50 centrifugation tube according to the manufacturers protocol (Stemcell Technologies). Residing erythrocytes were removed by using the Red Blood Cell Lysis Solution (Miltenyi Biotec). To isolate cells from tissues, tonsils and peripheral lymph nodes were cleaned, soaked in RPMI 1640 medium (5 % FCS, 10 mM HEPES; Gibco) and cut into fine pieces, whereas the colon was washed several times with cold PBS before the tissue was cut into small pieces (5 mm x 5 mm) in RPMI 1640. To remove the colonic epithelial cells, the colon pieces were incubated three times for 10 minutes at room temperature in epithelial buffer (DPBS, 5% FCS, 10 mM HEPES, 2 mM EDTA, pH 8.0; Gibco), shaken for 15 minutes at 37°C, and washed with PBS. The remaining colon tissue was cut into smaller pieces, digested with collagenase D (1 mg/ml; Roche), DNAse I (0.05 mg/ml; Roche) in RPMI 1640 (5% FCS, 10 mM HEPES) for 90 minutes in a shaker at 37°C. Tissue pieces from tonsils, lymph nodes and colon were further dissociated using a cell dissociation sieve-tissue grinder kit (Merck/ Sigma-Aldrich) and filtered through a cell strainer. The lymphocytes were then separated using a Lymphoprep gradient (Stemcell Technologies). To enrich the ILCs from the PBMCs or the tissue preparations the cells were either incubated with a lineage (lin) antibody cocktail (CD1a, CD3, CD19, TCRαβ, TCRγδ, CD14, CD11c, FcεRIα; all coupled to APC) and anti-APC microbeads (Miltenyi Biotec) or with antibodies and beads of the Human Lineage Cell Depletion Kit (Miltenyi Biotec). Washed cells were magnetically separated by applying the “deplete” program on the autoMACS Pro (Miltenyi Biotec). The lin⁻ cell fraction was subsequently used for ILC sorting. Magnetically sorted cells were stained with LIVE/DEAD Fixable Near-IR dye (Invitrogen) and ILC marker antibodies (lin^-^ cells) prior to flow cytometric cell sorting into NK cells (CD127^-^CD45^+^ population expressing either CD56 or NKG2A, or both), ILC1 (CD127^+^CD45^+^CD56^-^CD117^-^CRTH2^-^), ILC2 (CD127^+^CD45^+^CD56^-^CRTH2^+^) and ILC3 (CD127^+^CD45^+^CD56^-^CD117^+^CRTH2^-^NKp44^+^) using a FACSAria II (BD Biosciences). Cultured cells were first stained with a viability dye (LIVE/DEAD fixable near-IR dead cell dye, ThermoFischer Scientific) at 4°C for 15 min in the dark. Afterwards, cells were washed with staining buffer (0.5% BSA in PBS) and pelleted by centrifugation at 400×g for 5 min at 4°C. Single-cell suspensions were labelled with antibodies (see section above) for 15-30 min at 4°C, while chemokine receptor staining was performed at 37°C for 30 min. Cells were then washed and resuspended in staining buffer, followed by acquisition on a LSR-II SORP or LSRFortessa (BD Biosciences). For intracellular cytokine staining, ILCs were fixed and permeabilized using the Foxp3/Transcription Factor Staining Buffer Set (eBioscience) in accordance with the manufacturer’s protocol. Intracellular markers were labelled by overnight incubation with specific antibodies at 4°C. For cytokine detection, cells were stimulated in Yssel’s medium containing 100 ng/ml phorbol 12-myristate 13-acetate (PMA; Sigma-Aldrich) and 1 µg/ml ionomycin (Sigma-Aldrich) at 37°C. After 2 hours, Brefeldin A (5 µg/ml; eBioscience) was added and cells were incubated for an additional 4 hours to allow intracellular cytokine accumulation. Flow cytometric analysis was performed using a LRR-II SORP or LSRFortessa (BD Biosciences). FlowJo software, v10 (BD Biosciences), was used for data analysis.

### Isolation of gDNA and RNA for quantitative RT-PCR

Genomic DNA (gDNA) from primary or cultured cells was prepared using the DNeasy Blood and Tissue Kit according to the manufacturer’s protocol (Qiagen). The gDNA was then concentrated using the Genomic DNA Clean & Concentrator™-10 kit as recommended by the manufacturer (Zymo Research). The concentration of gDNA was measured using a Nanodrop photometer. The purified DNA was stored at -20°C until further use. Total RNA was extracted from primary or cultured cells using the RNeasy Plus Mini or Micro Kit (Qiagen) and quantified using a NanoDrop spectrophotometer (Thermo Fisher). cDNA was synthesized using the Transcriptor First Strand cDNA Synthesis Kit (Roche) or SuperScript IV VILO Master Mix (Thermo Fisher) with random hexamers. qRT-PCR was performed with SYBR Green Master Mix (Roche or Thermo Fisher) on a LightCycler 96 (Roche) or QuantStudio 3 (Applied Biosystems) using gene-specific primers for *HPGDS* (for: 5-ACCAGAGCCTAGCAATAGCAA-3, rev: 5-AGAGTGTCCACAATAGCATCAAC-3), *NOX2* (for: 5-ACCGGGTTTATGATATTCCACCT-3, rev: 5-GATTTCGACAGACTGGCAAGA-3) and *NRROS* (for: 5-AGTCTGCAAGTTGGTGGGTG-3, rev: 5-GGTCTTGAGAGGGTTGGCAT-3). Expression levels were normalized to *RPS9* (for: 5-CTTAGGCGCAGACGGGGAAGC-3, rev: 5-CGAAGGGTCTCCGCGGGGTCACAT-3) and calculated using the ΔCt or ΔΔCt method.

### Whole genome methylation analysis

Whole genome methylation analysis from ILC1, ILC2, ILC3 and NK cells of healthy adult donors was performed by Bi-seq using the Accel-NGS Methyl-Seq DNA Library Kit (Swift Biosciences) as recently described ^48^. Whole genome methylation analysis of ILC2 from juvenile, healthy donors or patients with atopy and asthma was performed by EM-seq using the NEBNext Enzymatic Methyl-seq Kit (New England Biolabs) according to the manufacturer’s protocol. Briefly, 50 ng genomic DNA in low TE buffer was sheared to ∼300 bp using a Covaris S2 (10% duty cycle, intensity 4, 200 cycles/burst, 80 s). Fragmented DNA was end-repaired and A-tailed (NEBNext Ultra II End Prep), ligated to EM-seq adaptors, and purified using magnetic beads. Oxidation was performed with TET2 and Fe(II) at 37°C for 1 h, followed by addition of stop solution and bead-based cleanup. DNA was denatured with formamide (85°C, 10 min) and subjected to APOBEC-mediated deamination at 37°C for 3 h. Libraries were purified, PCR-amplified (6 cycles) with Q5U Master Mix and indexed primers, and cleaned using magnetic beads. Finally, the libraries from Bi-seq or EM-seq were assessed using an Agilent Fragment Analyzer and sequenced on an Illumina NovaSeq 6000 with a depth of 206 million to 335 million paired-end reads (2 × 100 bp) or with a depth of 223 to 315 million paired-end reads (2 x 150 bp), respectively. Sequencing data were processed using the nf-core/methylseq pipeline (version 2.2.0), using default parameters, except that trimming was performed according to the Accel-NGS Methyl-Seq DNA Library Kit (and NEBNext Enzymatic Methyl-seq) manufacturer’s recommendation (--clip_r2 15 and --three_prime_clip_r1 15) and the relax_mismatches argument was used.^49^ Briefly, the pipeline employs FASTQC (version 0.11.9), trimgalore (version 0.6.7), and bismark (version 0.24.0) for read-level quality control, adapter trimming, bisulfite-aware alignment against the reference genome GRCh38.p13 (obtained from GENCODE.org), and cytosine-level DNA methylation quantification. Methylation calls were analyzed with the Bioconductor package BSseq (version bsseq_1.34.0 implemented in R 4.2.1)^50^. The raw methylation values of the samples were imported with the read.bismark function and smoothed using the BSmooth function to obtain estimated methylation value for every CpG based on a genome-wide methylation profile. Methylation profiles of CpGs with a minimum coverage of 2 mapped reads in at least two replicates of one condition, served as input for the detection of differentially methylated regions (DMRs) with the dmrFinder function (bsseq). The t-statistics were computed and DMRs identified using a quantile-based cutoff (0.01 and 0.99) and a maximum distance of 200 bp between each CpG to define a continuous DMR. DMRs were filtered to keep only those with a mean methylation difference (across the DMR) of at least 0.25 (25 %) between the two compared conditions incorporating at least three CpGs. Finally, DMRs were ranked by the sum of the t-statistics of each CpG associated (areaStat parameter) as suggested (bsseq documentation) and annotated with additional genomic information, including the nearest upstream/downstream gene and transcription start site. Annotation data were taken from GENCODE.org (Homo sapiens: GRCh38.p13; release 34) and corresponding Ensembl biomart dataset (Homo sapiens 100). The data evaluation was supplemented by hierarchical cluster analysis (pltree from cluster package v2.1.1), heatmap and PCA visualization (heatmap and prcomp from stats package, R Core Team). Differentially methylated regions (plotting BSmooth methylation estimates) were visualized with the plotRegion function from the bsseq package.

### *Ex vivo* culture and cytokine stimulation of human ILC2s

Sorted primary human ILC2s were cultured in V-bottom 96-well tissue culture plates in complete Yssel’s medium supplemented with cytokines and maintained at 37°C in a humidified incubator with 5% CO₂. Cells were washed with PBS containing 0.5% BSA and split every four days to maintain optimal density. To investigate context-dependent responses, cells were stimulated with defined cytokine cocktails (all purchased from Miltenyi Biotec) as follows: basal (IL-2, 10 ng/μl and IL-7, 10 ng/μl), type I (IL-1β, 20 ng/μl; IL-12, 20 ng/μl; IL-2, 10 ng/μl; IL-7, 10 ng/μl), type II (IL-2, 10 ng/μl; IL-7, 10 ng/μl; IL-33, 20 ng/μl; IL-25, 20 ng/μl; TSLP, 20 ng/μl), and type III (IL-1β, 20 ng/μl; IL-23, 20 ng/μl; TGF-β, 10 ng/μl; IL-2, 10 ng/μl; IL-7, 10 ng/μl). For cytokine switch experiments, cells cultured under type II conditions for 14 days were transitioned to type III cytokine media and maintained as indicated.

### Luciferase reporter assays

Differentially methylated regions (DMRs) and the corresponding promoter region from the *HPGDS* locus and *CRTH2* promoter region containing the enhancer element, as described by Quapp et al.^51^, were cloned into the pNL1.1-NanoLuc vector (Promega). To this end, PCR-amplified fragments from human PBMC-derived genomic DNA were digested alongside the luciferase vector with compatible restriction enzymes, gel-purified, ligated using T4 DNA ligase (NEB), and transformed into NEB 5-alpha *E. coli*. Following primers were used: *HPGDS*-promoter (for: 5-CGATGCTAGCAGCATGTTTACACTCC-3, rev: 5-GCATAAGCTTTGTATAACCAGCAGTCT-3), *HPGDS-*DMR (for: 5-GCATGGTA CCACATCCATCTGTATAATAAACT-3, rev: 5-ATGCGCTAGCGAGCAAATACCGC-3) and *CRTH2*-promoter (for: 5-CGATGGTACCTACCCCACTGTACTTTCCTGGC-3, rev: 5-GTCAGCTAGCAGAGAGGAAATAGCTGTCAGCCC-3). Positive clones were verified by sequencing. For the Luciferase assay, reporter constructs were nucleofected with the pGL4.53-PGK-Firefly control plasmid (Promega) into primary ILC2s using the Amaxa™ 4D-Nucleofector™ (program EH-110) and Nucleofector™ Solution (Lonza). Cells were cultured in Yssel’s medium supplemented with type II cytokines for 48 h. Luciferase activity was measured using the Nano-Glo Dual-Luciferase Reporter Assay System (Promega) according to the manufacturer’s protocol. Relative transcriptional activity was calculated as the NanoLuc/Firefly ratio and normalized to control.

### CRISPR/Cas9 knockout and HDR-mediated DMR deletion

In order to functionally delete specific DMR-associated genes, three guide (cr)RNAs for each DMR-associated gene were custom designed by IDT (Table S3). crRNA and tracrRNA (both 200 µM) were annealed at 95°C for 5 min and cooled to room temperature before complexing with recombinant Cas9 (IDT) to form ribonucleoproteins (RNPs). For RNP assembly, 18 μl of the duplex crRNA/tracrRNA and 7 μl of Cas9 reaction buffer (1X in water) were mixed, incubated at room temperature for 20 min, and immediately placed on ice. Equal amounts of RNP complexes targeting the same gene were mixed (e.g. 3 x 2μl/reaction). In vitro expanded ILC2 (1 x 10^6^ cells per sample) were mixed with 5 μl RNP mix and 3 μl electroporation enhancer (IDT), followed by incubation for 5 minutes at room temperature. Next, 100ul of P3 buffer (Lonza) was added, gently pipetted, transferred to a nucleofection cuvette and electroporated in an AmaxaTM 4D-Nucleofector device (program EH100). Finally, the cell solution was added to prewarmed Yssels medium. After 24 hours, the medium was replaced with Yssels medium supplemented with type II cytokines and cultured for 5 additional days before analysis.

CRISPR/Cas9-mediated deletion of differentially methylated regions (DMRs) was performed in expanded human ILC2s using synthetic guide RNAs (crRNAs) and custom-designed homology-directed repair templates (Table S3). RNP complex formation was performed as described above. crRNA activity was confirmed by in vitro cleavage of a 400-bp DMR-spanning PCR product, followed by proteinase K digestion and agarose gel electrophoresis. ILC2s (1 x 10⁶ cells per sample) were nucleofected with 4 µl RNP, 2 µl HDR Template (HDRT), and 3 µl electroporation enhancer in P3 buffer (Lonza) using the Amaxa 4D-Nucleofector (program EH115). Cells were cultured in Yssel’s medium with type 2 cytokines and 1 µM Nedisertib (M3814, Sigma-Aldrich) for 24 h, followed by cytokine-only medium. Cells were harvested on day 5 or 6 for analysis by qRT-PCR or flow cytometry. Editing efficiency was assessed by PCR and restriction digestion; transcript levels were analyzed by RT-PCR. Targeting of the Adeno-Associated Virus Integration Site 1 (AAVS1) served as a negative control.

### Pyrosequencing

Isolated gDNA (1-500 ng/reaction, according to the source) was first converted using the EZ DNA Methylation Lightning Kit (Zymo) according to the manufacturer’s protocol. The concentration of the converted DNA was determined using a NanoDrop photometer. Next, the region of interest was amplified by PCR using primer pairs (Table S1) and the ZymoTaq PreMix according to the manufacturer’s recommendations. The PCR amplicon was analyzed by agarose gel electrophoresis to confirm a correct amplification result. Pyrosequencing was performed using the PyroMark Q24 pyrosequencer (Qiagen) and PyroMark Q24 Advanced Reagents (Qiagen) as described previously.^21^ The sequencing data were analyzed by the PyroMark software according to the manufacturer’s recommendations.

### Motif Enrichment and TF Binding Site Prediction

De novo motif enrichment analysis was performed on DMRs (±20 bp) from ILC2 vs. ILC1, ILC3, and NK comparisons using MEME,^52^ applying the following parameters: -dna, -revcomp, -nmotifs 20, -mod anr. Motifs with E < 0.05 were matched to known transcription factor binding profiles using TOMTOM^53^ and the HOCOMOCO v11 core human database.^54^ Predicted TFs were filtered based on expression in human ILC subsets.^23^

### FIMO Analysis

FIMO (Find Individual Motif Occurrences) analysis^55^ was performed using motif sequences corresponding to core transcription factors identified in ILC2-specific DMRs. Motif position weight matrices were downloaded from the HOCOMOCO v11 core human database ^54^. DNA sequences from ILC2-specific DMRs (±20 bp) were used as input, with the following parameters applied: p-value < 1E-4, both strands, and default background model. FIMO output provided the exact binding motif sequences identified within the DMRs as well as their genomic coordinates. The predicted motif binding sites were parsed and mapped to the corresponding DMR regions.

### Gene-enrichment and functional annotation analysis

Identification of gene ontology (GO) terms or disease terms the online bioinformatic tool DAVID, v6.8, was used.^56^ Gene lists based on the DMR-associated genes were submitted with *Homo sapiens* as background. Enriched GO terms (biological process, molecular function, and cellular component) or Genetic Association Disease Database (GAD) terms were then identified by functional annotation clustering. Statistics based on *p*-value or Fisher’s exact value were generated by DAVID.

### Statistical analysis

Data were analyzed using GraphPad Prism. Normality test was assessed using the Shapiro-Wilk test, and appropriate parametric or non-parametric tests were applied accordingly. Two-tailed paired Student’s t-tests were used for paired comparisons. Linear mixed-effects analysis or one-way ANOVA (repeated-measures) with Dunnett’s multiple comparisons was used for multiple group comparisons. Data are presented as mean ± SEM. Significance thresholds were defined as follows: *p* > 0.05, ns; *p* ≤ 0.05, *; *p* ≤ 0.01, **; *p* ≤ 0.001, ***; *p* ≤ 0.0001, ****.

## Supplemental information

Document S1. Figures S1–S8, Tables S1-S3

