## Supplemental Data for "Human ILC methylomes uncover stable lineage codes and disease-linked ILC2 regulators in allergy"

### Supplementary Figure 1

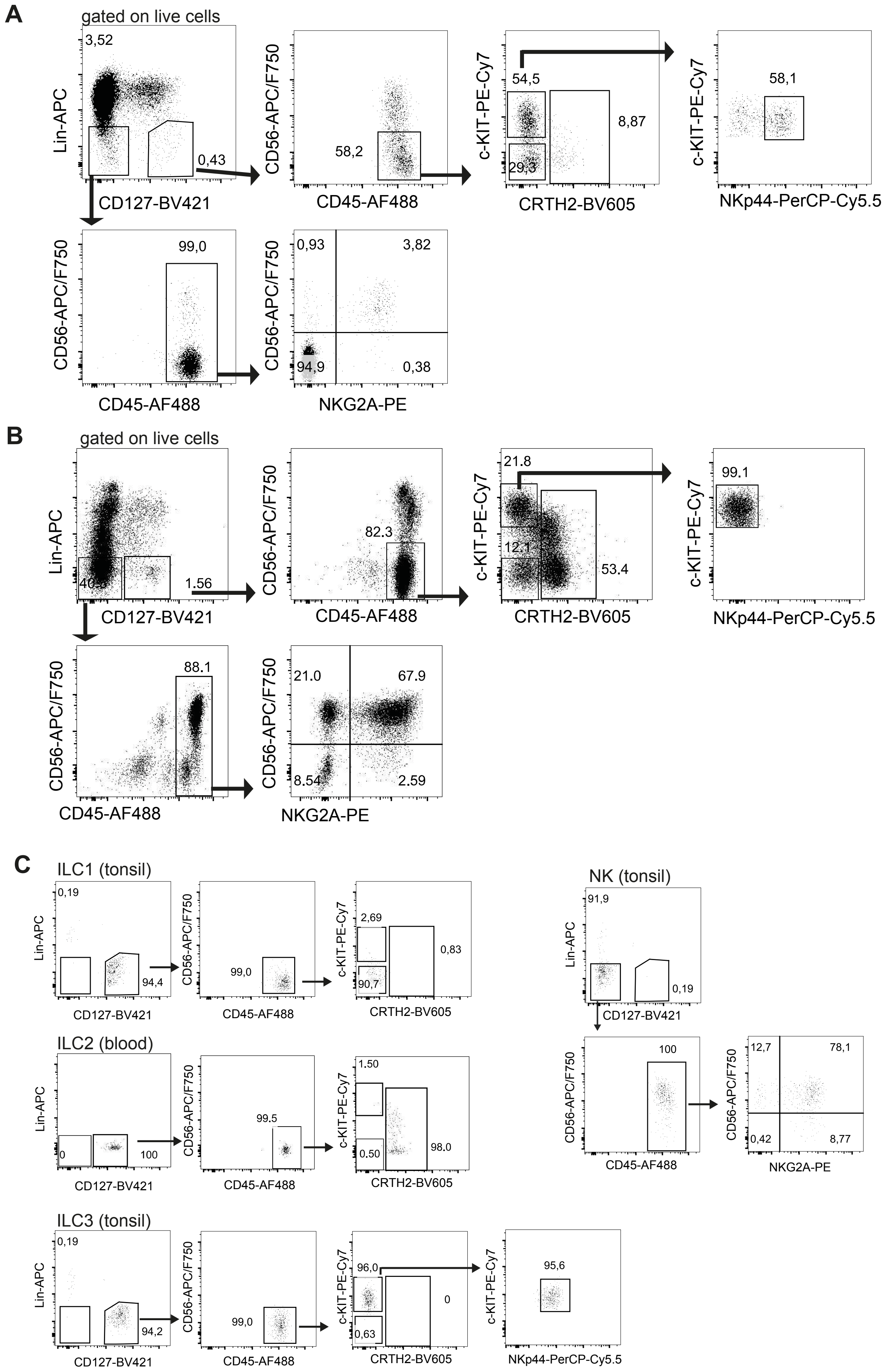

**Figure S1: Sorting strategy and purity control of human ILC populations for BS-seq.** Human ILCs from (**A**) non-inflamed tonsil and (**B**) peripheral blood were pooled from several donors and sorted by flow cytometry according to surface markers. ILC1 (lin$^-$CD127$^+$CD45$^+$CD56$^-$CD117$^-$CRTH2$^-$), ILC2 (lin$^-$CD127$^+$CD45$^+$CD56$^-$CRTH2$^+$), ILC3 (lin$^-$CD127$^+$CD45$^+$CD56$^-$CD117$^+$CRTH2$^-$NKp44$^+$) and NK cells (lin$^-$CD127$^-$CD45$^+$CD56$^+$NKG2A$^+$, lin$^-$CD127$^-$CD45$^+$CD56$^+$NKG2A$^-$, and lin$^-$CD127$^-$CD45$^+$CD56$^-$NKG2A$^+$). (**C**) Purity analysis of sorted NK cells, ILC 1, ILC2 and ILC3.

### Supplementary Figure 2

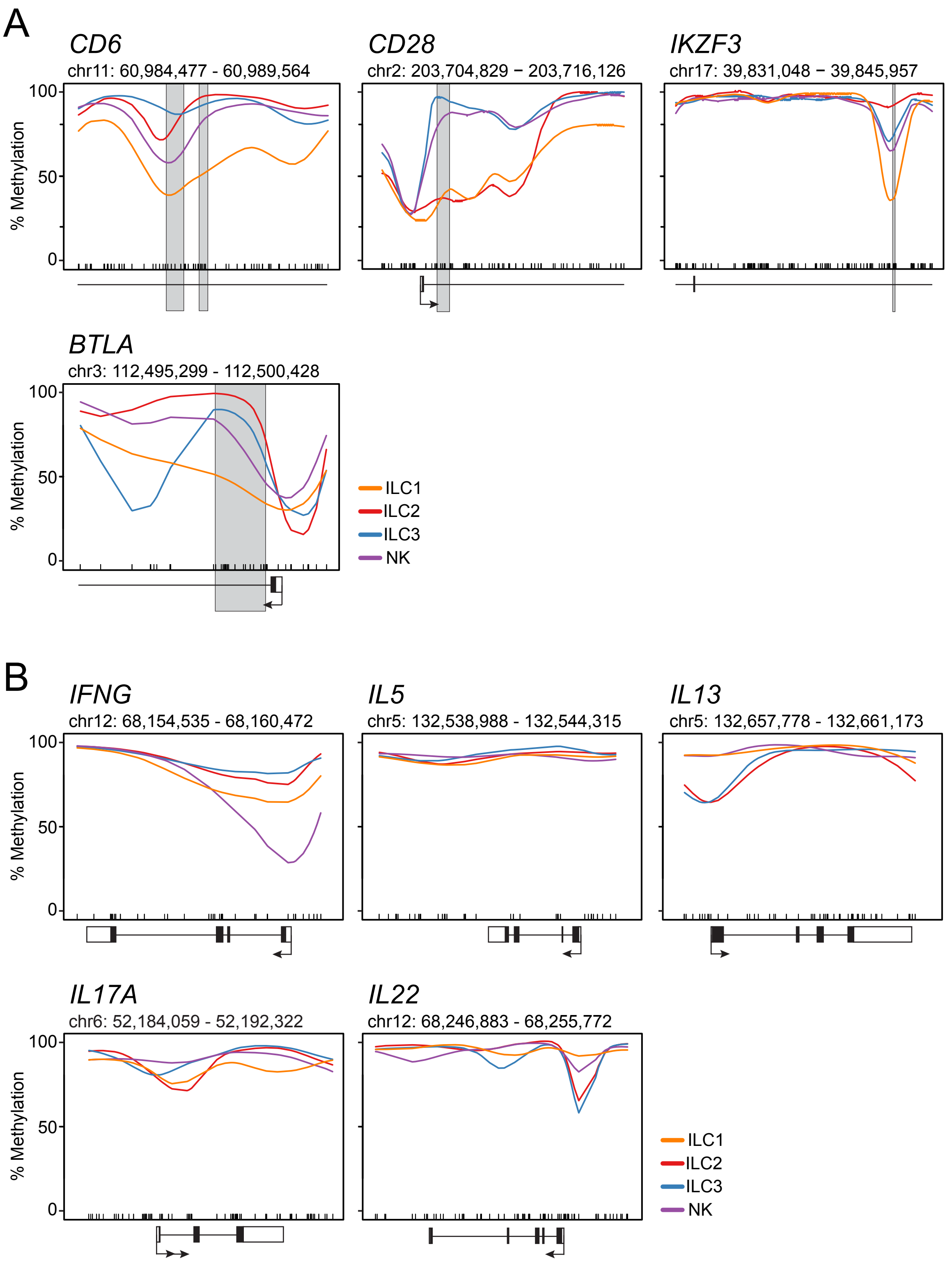

**Figure S2: Methylation line plots visualize the status of CpG motifs within gene loci associated with ILC1 and selected cytokines.** (**A**) Methylation line plots of ILC1 marker regions (*CD6*, *CD28*, *IKZF3* and *BTLA*). (**B**) Methylation line plots of selected Type I, -II and -III associated cytokine regions (*IFNG*, *IL5*, *IL13*, *IL17A* and *IL22*). The methylation line plots showing genomic methylation (0-100%) as a line within the specified chromosomal location in ILC1 (orange), ILC2 (red), ILC3 (blue), and NK cells (cyan). The position of the CpG motifs is indicated by a barcode. The gene elements transcription start site (arrow), translated (black box) and untranslated exon (white box), and the position of the DMR (grey box) are shown below or overlie the plot.

### Supplementary Figure 3

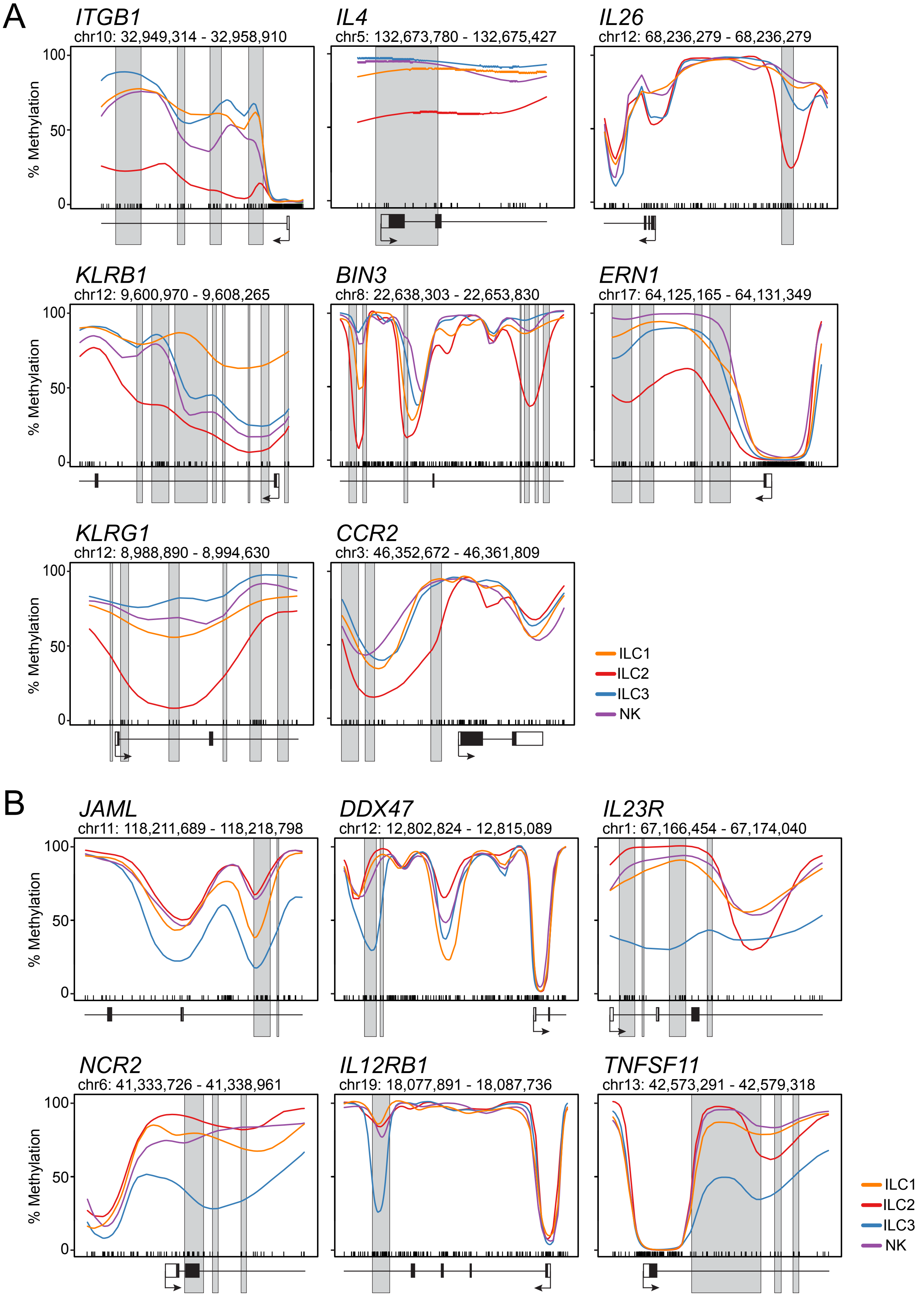

**Figure S3: Methylation line plots visualize the status of CpG motifs within gene loci associated with ILC2 and ILC3.** (**A**) Methylation line plots of ILC2 marker regions (*ITGB1*, *IL4*, *IL2*, *KLRB1*, *BIN3*, *ERN1*, *KLRG1* and *CCR2*). (**B**) Methylation line plots of ILC3 marker regions (*JAML*, *DDX47*, *IL23R*, *NCR2*, *IL12RB1* and *TNFSF11*). The methylation line plots showing genomic methylation (0-100%) as a line within the specified chromosomal location in ILC1 (orange), ILC2 (red), ILC3 (blue), and NK cells (cyan). The position of the CpG motifs is indicated by a barcode. The gene elements transcription start site (arrow), translated (black box) and untranslated exon (white box), and the position of the DMR (grey box) are shown below or overlay the plot.

### Supplementary Figure 4

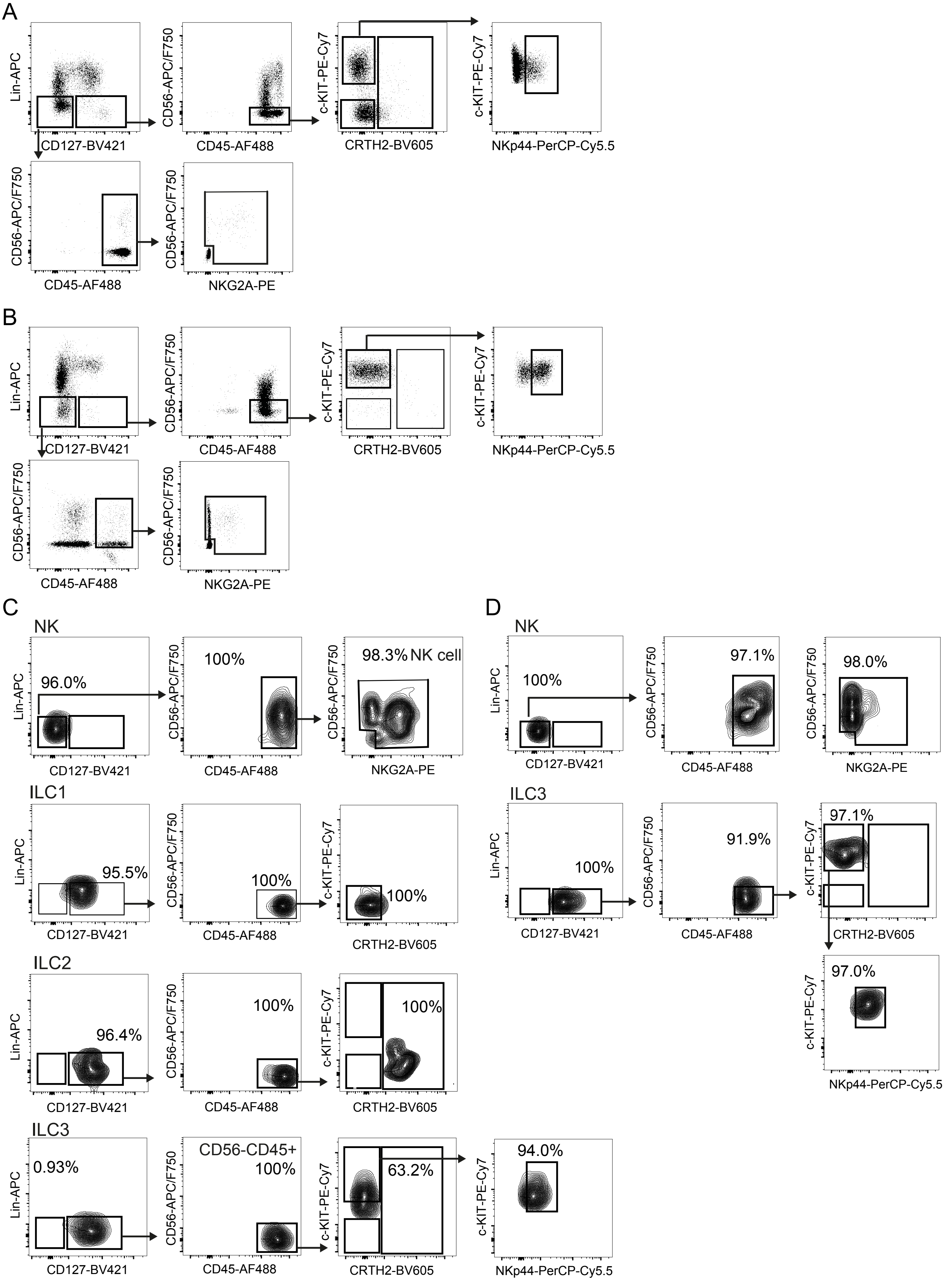

**Figure S4: Sorting strategy and purity control of ILCs from lymph nodes and colon.** Human ILCs from (**A**) non-inflamed LN and (**B**) non-inflamed colon were pooled from several donors and sorted by flow cytometry according to surface markers into ILC1 (lin$^-$CD127$^+$CD45$^+$CD56$^-$CD117$^-$CRTH2$^-$), ILC2 (lin$^-$CD127$^+$CD45$^+$CD56$^-$CRTH2$^+$), ILC3 (lin$^-$CD127$^+$CD45$^+$CD56$^-$CD117$^+$CRTH2$^-$NKp44$^+$) and NK cells (lin$^-$CD127$^-$CD45$^+$CD56$^+$NKG2A$^+$, lin$^-$CD127$^-$CD45$^+$CD56$^+$NKG2A$^-$, or lin$^-$CD127$^-$CD45$^+$CD56$^-$NKG2A$^+$). (**C**, **D**) Flow cytometry plots showing the purity analysis of sorted ILC subsets from LN (**C**) and colon (**D**).

### Supplementary Figure 5

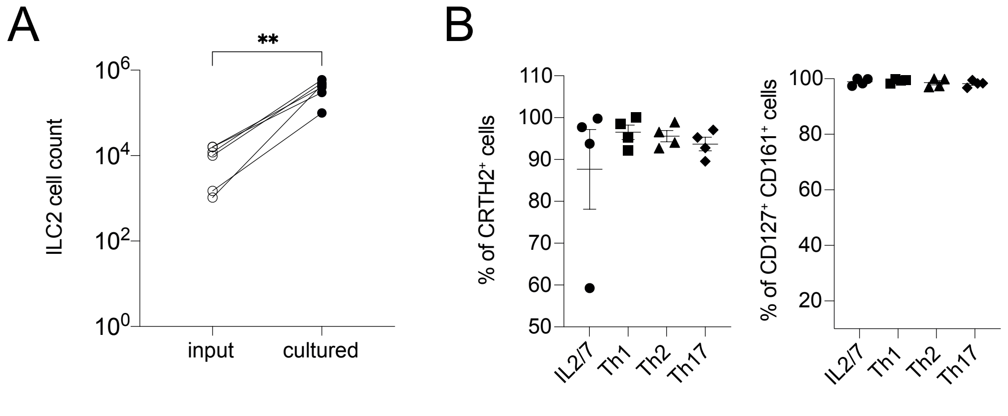

**Figure S5: *In vitro* expansion of ILC2.** ILC2 were isolated from the blood of healthy donors by flow cytometry and expanded in vitro under basal (IL-2/IL-7), type-I (Th1), type-II (Th2) or type-III (Th17) conditions. (**A**) Pairwise comparison of ILC2 counts, before (input) and after 2 weeks of culture with type-II cytokines (cultured). ($n = 6$, paired t-test, $p = 0.0029$). (**B**) Frequency of CRTH2$^+$ (left) and CD127$^+$CD161$^+$ ILC2 (right) after 2 weeks of culture under the indicated conditions.

### Supplementary Figure 6

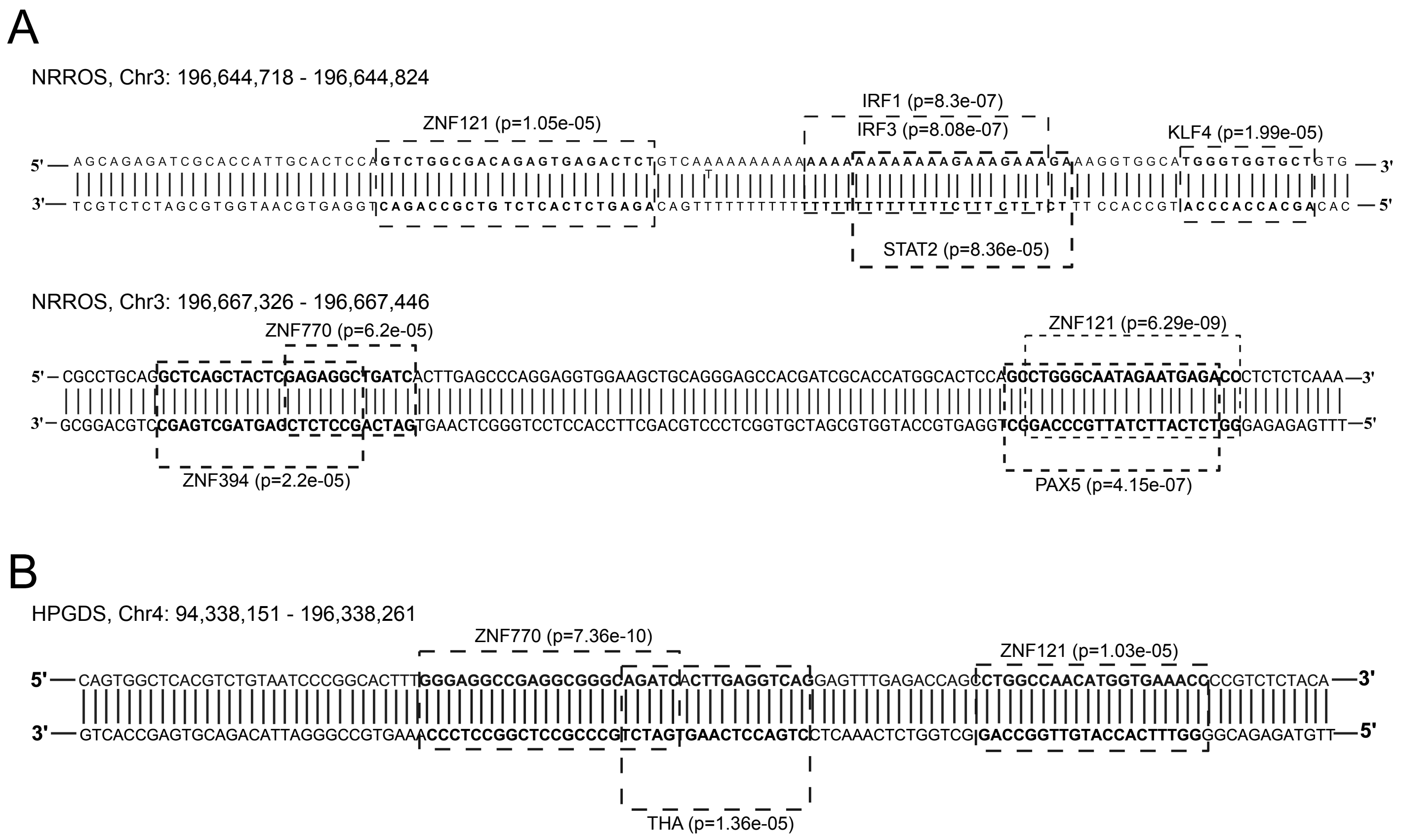

**Figure S6: Potential binding sites of core ILC2 TFs in NNROS and HPGDS DMRs.** Find individual motif occurrences (FIMO) analysis was performed using the DMRs that were identified in *NRROS* (**A**) and *HPGDS* (**B**). Binding sites of identified transcription factors are indicated by boxes along with their *p*-values.

### Supplementary Figure 7

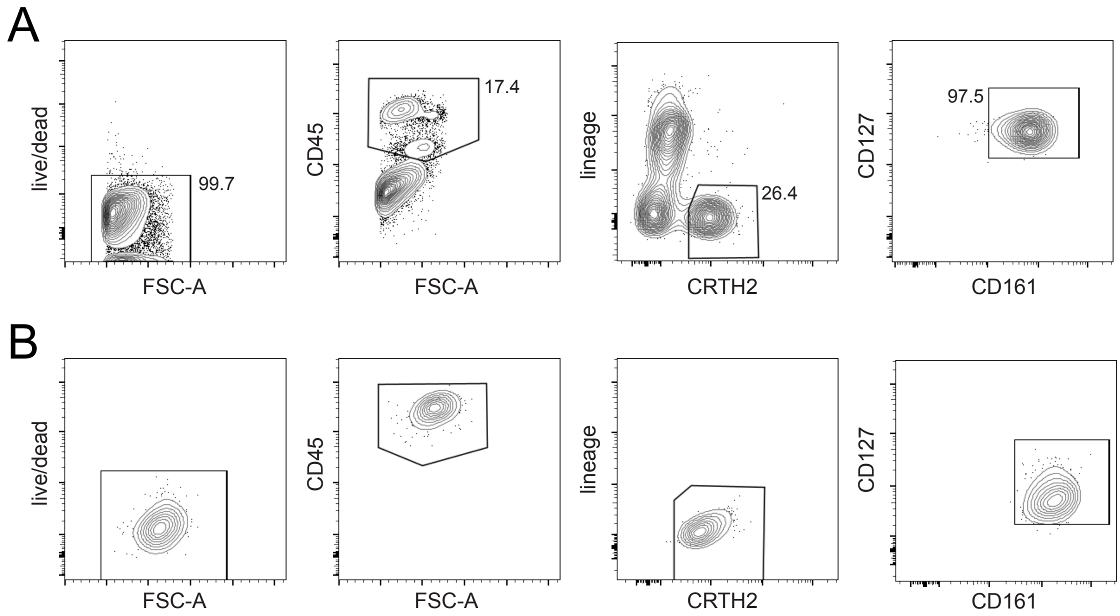

**Figure S7: Sorting strategy and purity control of ILC2 obtained from young donors.** Gating strategy for sorting of ILC2 from peripheral blood samples according to surface markers Lin$^{-}$CD45$^{+}$CD127$^{+}$CD161$^{+}$CD294$^{+}$ before (**A**) and after 14 days of in vitro expansion with type-II cytokines (**B**).

### Supplementary Figure 8

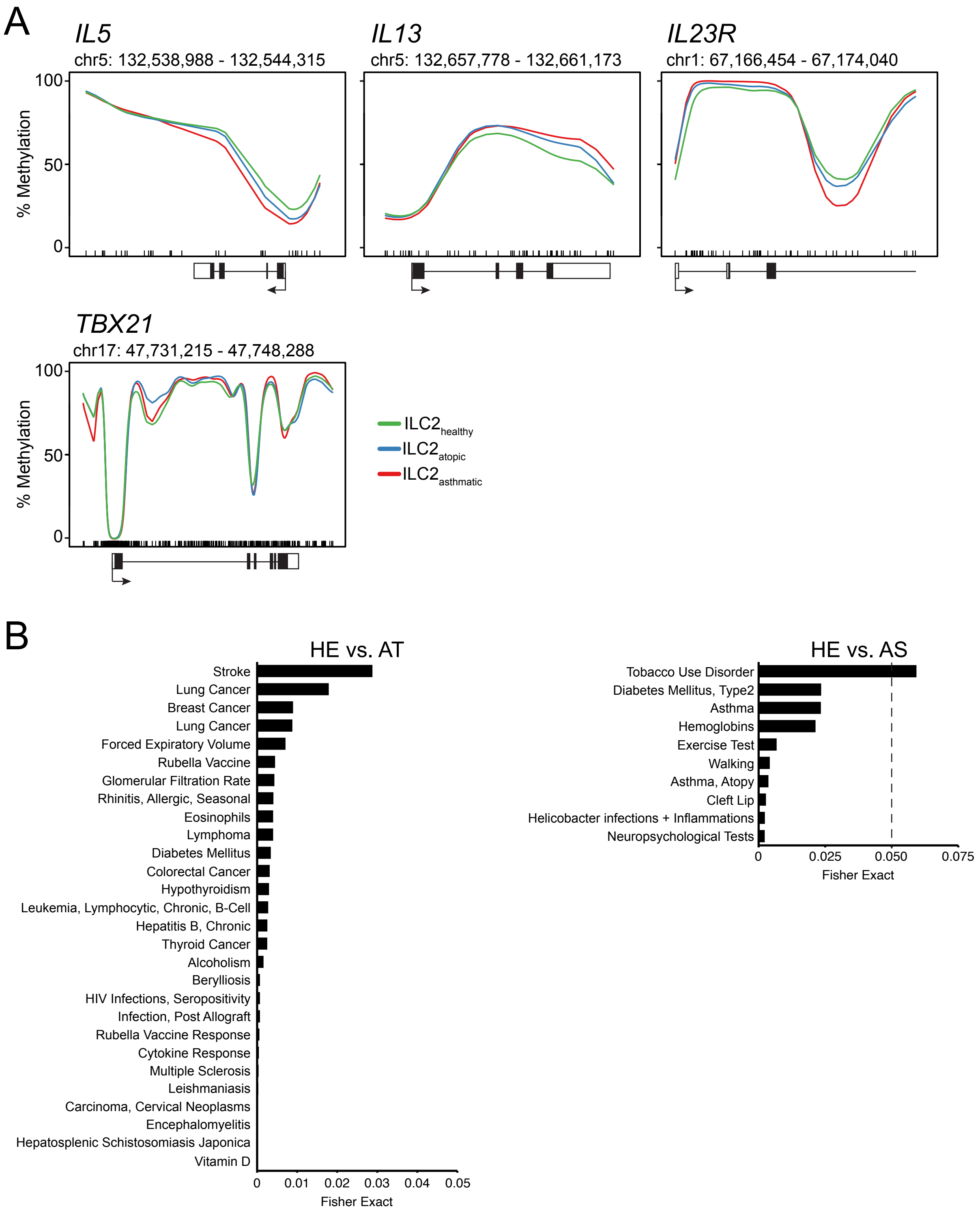

**Figure S8: Methylation line plots visualize the status of CpG motifs in selected gene loci in ILC2 from young donors.** (**A**) Methylation profiles of type I (*IL5*, *IL13*) and type III (*IL23R*, *TBX21*) associated gene loci in ILC2 from healthy, young donors, as well as atopic asthmatic patients after in vitro expansion in the presence of type-II cytokines. The methylation line plots showing genomic methylation (0-100%) as a line within the specified chromosomal location in ILC2 from healthy (green), atopic (blue), and asthmatic (red) donors. The position of the CpG motifs is indicated by a barcode. The gene elements transcription start site (arrow), translated (black box) and untranslated exon (white box), and the position of the DMR (grey box) are shown below or overlie the plot. (**B**) Functional annotation of disease-associated genes obtained from genes linked to DMRs, which were identified by pairwise methylome comparison of ILC2 from healthy donors and either atopic (HE vs. AT) or asthmatic donors (HE vs. AS). Disease classification is ranked according to Fisher's exact value.

**Table S1:** Localization of ILC marker regions, associated gene loci and amplification primer for pyrosequencing

| ILC population | Marker name | Marker location | | | DMR location | | | Associated gene | Ensembl gene ID | Pyrosequencing amplification primer forward | Pyrosequencing amplification primer reverse | Pyrosequencing primer 1 | Pyrosequencing primer 2 |
|---|---|---|---|---|---|---|---|---|---|---|---|---|---|
| | | Chromosome | Start | Stop | Chromosome | Start | Stop | | | | | | |
| ILC1 | PTPRC | 1 | 198678504 | 198678732 | 1 | 198678582 | 198678695 | PTPRC | ENSG00000081237 | AGATGGTTATGTTTTAATAAAAGTAAAGTT | CACCTCTCTACCTTTATAATCCTAATA | ACCTTTATAATCCTAATACCAA | ATAAAAACTAAACCCCAACA |
| | IKZF3 | 17 | 39843585 | 39843829 | 17 | 39843672 | 39843803 | IKZF3 | ENSG00000161405 | ATGTGATTTGATTATAGATGGTTTTTTTAT | ACCCTCTAACATTTAAATAACTCTCA | CTAACTACCTTAACATTCAT | |
| | CD28 | 2 | 203707728 | 203708011 | 2 | 203707404 | 203707954 | CD28 | ENSG00000178562 | AAGGATGTATAGGTGGTAGTTATAGGAG | ACCCCTATCAATCCAATTTTTAAACC | GTAGTTATAGGAGGTAGG | GATTTAGGTAAATTTAGGGTTAA |
| | LEF1 | 4 | 108140335 | 108140618 | 4 | 108163452 | 108163647 | LEF1 | ENSG00000138795 | AGATGGTTGATTATAAAATGGTATTGT | TATAAAACCCACTTCAACTCTACACTT | TGGGTTTATTTTTTATGTTGT | GGTTTTTGGGTGTAGTTA |
| | GRAP2 | 22 | 39920705 | 39920844 | 22 | 39927403 | 39927751 | GRAP2 | ENSG00000100351 | AGGTTTTGTTTTGTTAATGAGATAGTAAG | CTCCAACCCCCAAAATAAATTCCCTCACTC | AAAATCATAATATCTAAATATAACT | CATACCCTAACAACCAA |
| | BTLA | 3 | 112498215 | 112498495 | 3 | 112498113 | 112499161 | BTLA | ENSG00000186265 | TGGGTAGGATAGATGTTGGATAATT | AAACCTATAAATAACTCATCAAAAACTTC | GGATAATTGGGGTTATTTAATAT | ATTAGGGAGGAAGAAGAAT |
| | CD6_1 | 11 | 60986200 | 60986467 | 11 | 60986253 | 60986606 | CD6 | ENSG00000013725 | GAAGGAATGTAAAGAGAGAAAAGAAGT | AACCACACCCATTTTACCTC | AGAGAAAAGAAGTTTTAGTTAG | |
| | CD6_2 | 11 | 60986752 | 60987039 | 11 | 60986945 | 60987096 | CD6 | ENSG00000013725 | GGAGGTAGTTTTTGTTGTAGGTATATTTGT | TCCCTTACCTAAACCTCCTAAATAACC | GGTTAGGGTAATAGAAATG | GAGGTTAGGAGTTTAAGA |
| | CLEC2D | 12 | 9675143 | 9675350 | 12 | 9674926 | 9675487 | CLEC2D | ENSG00000069493 | GGTGGTAGGTGTTTGTAGTTT | TCTACTACTCTACTTTCTATCCTTTTTC | ATGGTGGGTGAATTT | |
| | TBX21 | 17 | 47734825 | 47735149 | 17 | 47741811 | 47741990 | TBX21 | ENSG00000073861 | GTGGGGAAGGGGTGTTTTAT | ACCCCACCTTTAATATCCACACCT | GGTTTAGGGTTTTTGAGT | TGGTATTTGTTTAGTAAGTAATT |
| ILC2 | NRROS_1 | 3 | 196644618 | 196644767 | 3 | 196644275 | 196644729 | NRROS | ENSG00000174004 | GTGTGATGGTGATTTGTAGTGA | CAAAATCTCACTCTATCCCCAAACTAAAATAC | GGTAGGAGAATTTTTTGAAT | |
| | NRROS_2 | 3 | 196667475 | 196667734 | 3 | 196667327 | 196667657 | NRROS | ENSG00000174004 | ATTTGTTTAAGAATTGTGGTGAAGATGTTA | CCAAACTATCACTACATCATACTACTACCT | ATCTTCCATCACTATATACTT | |
| | KLRB1 | 12 | 9603459 | 9603671 | 12 | 9603309 | 9603852 | KLRB1 | ENSG00000111796 | TTGAGGTTGGGAGTTAGAGATTAGTTTGATTA | TTCACTCTTATTACCCAAACTCATAATACAATCT | CATAATACAATCTCAACTCAC | |
| | ITGB1 | 10 | 32950759 | 32951073 | 10 | 32950853 | 32951944 | ITGB1 | ENSG00000150093 | AGTTTGGGTTAAGAAGTTTTAATAATAAAGG | TTAAATTTCCAAACTTTACCTCCCCCTAC | GGTTTATAAAATAATTTGAAATTTGAG | AGGGAAAGATAAATAATGGATATA |
| | MAF | 16 | 79592705 | 79593059 | 16 | 79592870 | 79593292 | MAF | ENSG00000178573 | AAGATTGAGATAGGTTGGAAGATTAA | ATTCATTCAACATCCTAACAAATAATC | ATTAAAATATAGGAGTATATGAGTG | ATTTTGTTGATTTGTAAGGT |
| | HPGDS | 4 | 94338134 | 94338240 | 4 | 94338146 | 94338404 | HPGDS | ENSG00000163106 | AAAAATTAGGTAGGGTGTAGTGGT | TATTAACCAAACTAATCTCAAACTCCTAAC | GGTAGGGTGTAGTGGTTTA | |
| | ERN1_1 | 17 | 64126432 | 64126690 | 17 | 64126172 | 64126540 | ERN1 | ENSG00000178607 | GTTTATTGAGTTTGTTATTAGGGTTGTTG | CCCTCAAACCTATACAAATAACACTA | TTTAGAGTATAGGAGAAAAATG | |
| | ERN1_2 | 17 | 64125544 | 64125905 | 17 | 64125408 | 64125945 | ERN1 | ENSG00000178607 | AGTGGTGTGATTTTAGTTTATTGTAATT | TCCCCACAAAACATCTTCTATAACCC | CCTTAATAAACCATCATCC | CCTATAATCCCAACACTTT |
| | IL26 | 12 | 68233918 | 68234079 | 12 | 68233460 | 68234175 | IL26 | ENSG00000111536 | AGGGTATTTTGTGGTGATGG | ATCTCCCTTTACTAAAATCTAAACTCT | ATGGTAGGGAGGTGT | GATTAGTTATTGTTTGGAATTG |
| | KLRG1 | 12 | 8957723 | 8958113 | 12 | 8957357 | 8958323 | KLRG1 | ENSG00000139187 | GGGATATTAAAGGGTAGGGAGTGT | CCAACCTAAATACTATAACAAAAACCTAT | TAAAGGGTAGGGAGTGTT | GATAAGGGTGGGGAA |
| | GATA3_1 | 10 | 8058853 | 8059074 | 10 | 8068329 | 8068827 | GATA3 | ENSG00000107485 | GTTTATTGGGGGGTTTTAAATTAGAT | CTCTTCCCTAAAACCTTTTCTCC | GGGGGTTTTAAATTAGATTTTAT | |
| | GATA3_2 | 10 | 8067703 | 8067900 | 10 | 8067476 | 8067814 | GATA3 | ENSG00000107485 | AGTTATTTGGGAGGTTGAGGT | ACAAAACAAAAACTATAAAAATCCTAATAC | GGAGAATGGTGTGAATTTA | |
| | IL10RA | 11 | 117991133 | 117991307 | 11 | 117988494 | 117988951 | IL10RA | ENSG00000110324 | GGTTGTAGTGAGTTAAGATTAAGTTATTG | ACACTATACCTCATAAACATATAAAATTATTACT | GTTATTGTATTTTAGTTTGGGT | |
| | CCR2 | 3 | 46356241 | 46356584 | 3 | 46356367 | 46356791 | CCR2 | ENSG00000121807 | AGGGAGTAGGAATTATGAGTTTAGAT | TCTCAACTTTCTACCTCCATCATCC | GTTTGGGGAATTTTAAGG | |
| | IL4 | 5 | 132673855 | 132674072 | 5 | 132673940 | 132674480 | IL4 | ENSG00000113520 | GTAGTTGGGAGGTGAGATTTAT | ATACTAATTAACCCCAAATAACTAACAA | GGGAGGTGAGATTTATTAA | AGAAGTTAAAGATGTAATGTTGATAAA |
| | BIN3 | 8 | 22651075 | 22651409 | 8 | 22651148 | 22651384 | BIN3 | ENSG00000147439 | AGGGATAGTTGGTGTTGAG | ATACCACTCCCAAATTCCTAAC | AGTTGGTGTTGAGAG | ATGTAAAGGAGAGAATATTTGA |
| ILC3 | IL23R | 1 | 67168817 | 67169131 | 1 | 67166878 | 67167381 | IL23R | ENSG00000162594 | AGTTTTTAGGTTGATTTTAGTTTTGTGTTA | TCTCATTCTATCACCCAAACTACAATACA | ATGGTGTTGAATATTAATAAAAG | AGTTTGGTTAATATGGTGAAA |
| | NCR1 | 19 | 54906788 | 54907076 | 19 | 54906910 | 54907314 | NCR1 | ENSG00000189430 | GTTGGATTTGGTGGTAATAGGTAA | ACATTCCTCAATCATACCCCTACATA | GTTTTTTAATTGGAGAGTGA | ATTTTTGTTTATAAGGGGTT |
| | IL12RB1 | 19 | 18079079 | 18079310 | 19 | 18079135 | 18079770 | IL12RB1 | ENSG00000096996 | TAAGATTATATATGTAAGTGGGATGGG | CAAACTCTACCTCCTAAATTCACAC | CTCCCCAATAACTAAAAC | ACTAAATTTTTATATTTTTAATA |
| | KIT_1 | 4 | 54661689 | 54662003 | 4 | 54661460 | 54662026 | KIT | ENSG00000157404 | ATGTGTAGTTTTGGTAATTTGAATAGA | ATCATATAAACCAAAAAAAACATATCTCC | TTGGTAATTTGAATAGATATTGT | AAGATTTTTAATAAGTAAAGAGTAG |
| | KIT_2 | 4 | 54663171 | 54663444 | 4 | 54662635 | 54662874 | KIT | ENSG00000157404 | AGGTTTAAGTTAAATTATTATTTAGGATAT | ACCTAATAATAAAATCCACACACTTC | ATTATTTAGGATATGTGGTATTA | ATGGAGATGATGATATAGG |
| | JAML | 11 | 118222767 | 118223076 | 11 | 118217075 | 118217579 | JAML | ENSG00000160593 | GTTATTTAGGTTGGTATGTAGTGGTATA | CTCAAAAAATTTTAACTATTAACCTCACCT | TTGGTATGTAGTGGTATAAT | AATTTTTGTATTTTTAGTAGAGA |
| | DDX47 | 12 | 12804375 | 12804689 | 12 | 12803937 | 12804598 | DDX47 | ENSG00000213782 | AGTTTGAGATATTAGATTTTTTTGGGAGTA | AAATCCCCAAAACCTATTTATTTACT | TTAGTGGGTTTAGGAG | ATTTTAGTTTTTTTGTGTTGT |
| | NCR2 | 6 | 41343265 | 41343580 | 6 | 41343437 | 41343942 | NCR2 | ENSG00000096264 | TTGGTTTTGAGGGTTGTTTAATT | TAATAACATTCCCTCTTCCCTACCTC | TTTTTTGTTTTAGAGATAAGGT | AGTGGTGGTATTATAGG |
| | TNFSF11_1 | 13 | 42575881 | 42576042 | 13 | 42575554 | 42577400 | TNFSF11 | ENSG00000120659 | TTGGTTTTATGTTTATAGTAAGTGGATATT | AATAACTTACCCATTTTCTCATCC | AGTAAGTGGATATTGTTTG | |
| | TNFSF11_2 | 13 | 42577453 | 42577453 | 13 | 42575554 | 42577400 | TNFSF11 | ENSG00000120659 | TGGAGAAAAGAAGGTATGTAAGT | ACCACACACTATTTTCTCTTTAACAA | AAAAGAATGTTTAGAATAGTGTAA | |
| | AHR | 7 | 17307659 | 17307986 | 7 | 17307560 | 17308047 | AHR | ENSG00000106546 | TTGAAGGAGTTTAATTAGAGGAGTGATAT | ACTAATTTAAACAACCTAAAAAAATCTCTC | ATAGTTAGATTTATGTTTTAGAAAG | TTAGTTTAATATTTAAGAATAGATG |
| | CXCR4 | 2 | 136115419 | 136115804 | 2 | 136115419 | 136115634 | CXCR4 | ENSG00000121966 | GGGAATAGTTAGTAGGAGGGTAGGGATTTA | CCCACCATCTACTCCATCATCTTCTTA | TGGTTTTTGATTGTTGG | TGTTTTGTATAGGAAGTTTTTAAA |
| | RORC | 1 | 151823213 | 151823504 | 1 | 151824462 | 151824497 | RORC | ENSG00000143365 | GAGGGAAGTAGGGTGGTTTTGATAA | CCTTTCTAACCCACTATTCCT | GGTGGTTTTGATAAGGAT | AGTTTAGATATTTTTTTTAAGGG |

**Table S2:** Clinical information about healthy, atopic and asthmatic juvenile donors of ILC2

| Sample name | Sample Group | Gender | Age | IgE | Atopy | Allergens with sIgE ≥0.7 kU/L | Blood eosinophils (cells/µl) | Treatment (inhalative corticosteroids) |
|---|---|---|---|---|---|---|---|---|
| HE1 | healthy | m | 12 | 21.1 | neg | na | 140 | na |
| HE2 | healthy | m | 15 | 15.7 | neg | na | 170 | na |
| HE3 | healthy | m | 14 | 117 | neg | na | 110 | na |
| HE4 | healthy | m | 11 | 38.9 | neg | na | 250 | na |
| AT1 | atopic | m | 16 | 1382 | pos | cat, grass, birch, house dust mite, haselnut, apple, | 140 | na |
| AT2 | atopic | m | 13 | 594 | pos | grass, plantain, apple | 80 | na |
| AT5 | atopic | m | 12 | 4100 | pos | grass, birch, house dust mite, haselnut, soy bean, apple | 640 | na |
| AT6 | atopic | m | 13 | 795 | pos | grass, birch, horse, haselnut, apple | 270 | na |
| AS1 | asthmatic | m | 12 | 537 | pos | dog, grass, house dust mite | 430 | Salmeterol/ Fluticason |
| AS2 | asthmatic | m | 10 | 1283 | pos | mugwort, grass, birch, house dust mite, peanut, walnut, sesame, tomato, apple | 710 | Salmeterol/ Fluticason |
| AS3 | asthmatic | m | 10 | 853 | pos | cat, grass, birch, house dust mite | 910 | Fluticason |
| AS4 | asthmatic | m | 15 | 440 | pos | alternaria alternata, Aspegillus fumigatus | 50 | Vilanterol/ Fluticason |

**Table S3:** CrRNAs and HDR templates for CRISPR/Cas9 mediated genome manipulation

| | | |
|---|---|---|
| **a) crRNAs used for CRISPR/Cas9-mediated knock-out** | | |
| **Target gene** | **crRNA** | **Sequence** |
| *HPGDS* | Hs.Cas9.HPGDS.1.AA | ATGTCATGTTGATGCTATTG PAM (TGG) |
| *HPGDS* | Hs.Cas9.HPGDS.1.AB | ATCCCCATTTTGGAAGTTGA PAM (TGG) |
| *HPGDS* | Hs.Cas9.HPGDS.1.AC | TCCAAGTCTTGCATAAGATG PAM (AGG) |
| *NRROS* | Hs.Cas9.NRROS.1.AA | GTACAACACCTCGTCGCCGA PAM (GGG) |
| *NRROS* | Hs.Cas9.NRROS.1.AB | CAACGTCAGCTACAACGTCC PAM (TGG) |
| *NRROS* | Hs.Cas9.NRROS.1.AC | GGGTCCGCAACTTGCTGTAC PAM (TGG) |
| **b) crRNA and HDR templates used for DMR knock-out** | | |
| **Target** | **crRNA (sequence)** | **HDR template (sequence)** |
| HPGDS DMR | crRNA 1 (- strand): TGAGCAAATACCGCATTACA (PAM: AGG) | 5–CAGTCTCAGAACCATTTAGTCCTCACTCTATGTCATGTCTGTCCTATATATGAGGTTCTGTTTTAA GACAATGCACCTCTAAGCTTTCTGATATAACATCTGGTATTCCTGAGACAAACAGAAAATATAATTAT GCAAATAACTACAGAAATGTCATGTAAAAAGT–3 |
| | crRNA 2 (+ strand: GTAATGCGGTATTTGCTCAG (PAM: AGG) | 5–ACTTTTTACATGACATTTCTGTAGTTATTTGCATAATTATATTTTCTGTTTGTCTCAGGAATACCAG ATGTTATATCAGAAAGCTTAGAGGTGCATTGTCTTAAAACAGAACCTCATATATAGGACAGACATGAC ATAGAGTGAGGACTAAATGGTTCTGAGACTG–3 |
| NRROS DMR1 | crRNA 1 (- strand): ACCGCATGGCACAGCTACTT (PAM: GGG) | 5–TCCACAGAAGGCCATGGTCCCATGATACACTCAGGGGTGCTGTTCAGAATATCAGGCACC GCATGGCACAGCTACTTGGGAAGCTTCCCTAGGTGGGAGGACTACTTGAGGCTGGGAGCTT GAGGCTGTAGTGTGCTACAATCCCATCTGTGAATAACTACTGCAC–3 |
| | crRNA 2 (+ strand): TCAGAATATCAGGCACCGCA (PAM: TGG) | 5–GTGCAGTAGTTATTCACAGATGGGATTGTAGCACACTACAGCCTCAAGCTCCCAGCCTCAA GTAGTCCTCCCACCTAGGGAAGCTTCCCAAGTAGCTGTGCCATGCGGTGCCTGATATTCTGA ACAGCACCCCTGAGTGTATCATGGGACCATGGCCTTCTGTGGA–3 |
| NRROS DMR2 | crRNA 1 (- strand): CGCTTAAGATTTGTTCAAGG (PAM: TGG) | 5–GTGCTAAGTGCTTTATGTGGATAATTTCGCTTAATCTTGGAGAAAACTCTAGGCTGAGAGC GCTTAAGATTTGTTCAAGGAAGCTTCAATTATAGATGTCATTTCTCTGGGCCTAGGCAAAGA CTGGGCACAGTGGCTCATGCCTGTAATCCCAGCACTTTGGGAG–3 |
| | crRNA 2 (+ strand): TGTTCAAGGTGGCTGGGCGT (PAM: GGG) | 5–CTCCCAAAGTGCTGGGATTACAGGCATGAGCCACTGTGCCCAGTCTTTGCCTAGGCCCAGA GAAATGACATCTATAATTGAAGCTTCCTTGAACAAATCTTAAGCGCTCTCAGCCTAGAGTTTT CTCCAAGATTAAGCGAAATTATCCACATAAAGCACTTAGCAC–3 |
